# Depletion of lamins B1 and B2 alters chromatin mobility and induces differential gene expression by a mesoscale-motion dependent mechanism

**DOI:** 10.1101/2023.06.26.546573

**Authors:** Emily M. Pujadas, Xiaolong Wei, Nicolas Acosta, Lucas Carter, Jiekun Yang, Luay Almassalha, Ali Daneshkhah, Suhas S.P. Rao, Vasundhara Agrawal, Fidan Seker-Polat, Erez Lieberman Aiden, Masato T. Kanemaki, Vadim Backman, Mazhar Adli

## Abstract

**BACKGROUND:** B-type lamins are critical nuclear envelope proteins that interact with the 3D genomic architecture. However, identifying the direct roles of B-lamins on dynamic genome organization has been challenging as their joint depletion severely impacts cell viability. To overcome this, we engineered mammalian cells to rapidly and completely degrade endogenous B-type lamins using Auxin-inducible degron (AID) technology.

**RESULTS:** Paired with a suite of novel technologies, live-cell Dual Partial Wave Spectroscopic (Dual-PWS) microscopy, *in situ* Hi-C, and CRISPR-Sirius, we demonstrate that lamin B1 and lamin B2 depletion transforms chromatin mobility, heterochromatin positioning, gene expression, and loci-positioning with minimal disruption to mesoscale chromatin folding. Using the AID system, we show that the disruption of B-lamins alters gene expression both within and outside lamin associated domains, with distinct mechanistic patterns depending on their localization. Critically, we demonstrate that chromatin dynamics, positioning of constitutive and facultative heterochromatic markers, and chromosome positioning near the nuclear periphery are significantly altered, indicating that the mechanism of action of B-type lamins is derived from their role in maintaining chromatin dynamics and spatial positioning.

**CONCLUSIONS:** Our findings suggest that the mechanistic role of B-type lamins is stabilization of heterochromatin and chromosomal positioning along the nuclear periphery. We conclude that degrading lamin B1 and lamin B2 has several functional consequences related to both structural disease and cancer.

## BACKGROUND

Chromatin is the complex macromolecular polymer-assembly formed by folding the genome and its associated proteins within the confines of the cell nucleus. Numerous studies indicate that the interplay of physical, chemical, and molecular principles leads to the emergence of the major organizational features of the genome, including polymer-polymer interactions, chromatin loop extrusion^1,2^, phase separation^3,4^, the physical interaction of chromatin with stable architectural elements of the nucleus (i.e., nuclear envelope)^5–7^, and chromatin dynamics^8,9^. Our understanding is limited, however, about the mechanisms governing chromatin folding due to the tethering that occurs at the nuclear periphery.

It is widely believed that the nuclear lamina is the structural limit on the spatial distribution of 3D genome organization in the nucleus^10^. Lamin proteins are structural components of the nuclear lamina that interact with both the cytoskeleton and the genome^11^. Lamins interact with chromatin either indirectly or directly through chromatin binding proteins^12^. Mechanistically, these lamin-proteins are also involved in several critical processes, including transcription, DNA repair, and replication. Lamin dysregulation is associated with more than 15 diseases, termed laminopathies^11,13,14^. The four main lamin isoforms found in mammals are lamin A/C, lamin B1, and lamin B2, respectively^11^. Although lamin B1, lamin B2, and lamin A/C form overlapping networks, B-type lamins localize primarily to the nuclear periphery adjacent to the inner nuclear membrane, while A-type lamins can extend into the nucleoplasm^6,15^. In contrast to A-type lamins, B-type lamins remain tightly associated with the nuclear membrane and have been mapped using DamID to reveal the existence of dynamic and functional euchromatin lamin B1 domains.^16^ Prior work identified that inhibition of lamin A/C and lamin-B receptor (LBR) alters A/B compartment segregation, specifically chromatin region localization^17,18^. Differential inhibition of lamin B1 or B2, in contrast, has been shown to have variable effects on compartmentalization. These varied effects are hypothesized to be due to the persistence of the conjugate B-protein.^6,14,19^ As B-type lamins have similar domain structures, lamin B1 and lamin B2 were conjectured to have redundant functions^20^. It has also been postulated that silencing B-type lamin expression leads to a dramatic increase of lamin A mobility in the nucleoplasm to maintain chromatin organization and transcription.^21^ These varied hypotheses illustrate a need to uncover how lamins B1 and B2 contribute to normal cellular physiology.

Mechanistically, B-type lamins are thought to regulate chromatin structure and gene transcription by physical attachment of chromatin to constrain chromatin dynamics and structure^6^. This mechanism has been evidenced by the recruitment of genes to the nuclear lamina within regions termed lamin-associated domains (LADs), associated with transcriptional repression, B-compartments, and heterochromatin nucleosome markers (e.g., H3K9me2/3)^16,22–24^. Paradoxically, transcriptionally rich territories exist adjacent, and interspersed within, these sites of transcriptional repression, suggesting that the nuclear lamina forms a complex environment for transcriptional regulation^25^. Physically, LADs are variably sized, with domains spanning from 100 kb to 10 Mb and a median size of 0.5 to 1 Mb from ChIP-seq analysis of lamin A/C and lamin B1.^24^ However, the reported relative sizes of LADs can also vary between molecular and imaging techniques or cell type. For example, while imaging studies in mouse and human cells report LADs to be 10 kb to 10 Mb in size, ChIP-seq studies in *Drosophila* cells report LADs to vary between 7 and 700 kb^23,26,27^.

Despite the broad role of B-lamins in physiology, the essentiality and redundancy of B-lamin proteins poses a formidable challenge to understanding their distinct role in regulating cell function. To overcome this limitation, we generated an auxin-inducible degron system targeting both B-type lamins simultaneously (see **Materials and Methods).** Utilizing this engineered cell line and based on prior studies of lamin A/C compared to lamin-B receptor in nuclear inversion^17,18^, we tested the hypothesis that the mechanism of action of B-type lamins is due to their role in tethering the genome to the nuclear periphery. In the predominant model of lamin B1/B2 at present, these lamins act by the differential constraint of inner nuclear matrix to regulate chromatin higher-order structure and dynamics^5,6,16,23^. Based on these prior studies, we hypothesized that simultaneous LMNB1 and LMNB2 inhibition would result in (1) increased chromatin chain fluctuations, (2) internal translocation of heterochromatin domains, (3) dissociation of genes from the nuclear periphery, and (4) that the genes normally located on the nuclear periphery would be differentially expressed due to the transformation in mesoscale (∼10 – 1000 nm)^28–30^ chromatin folding.

Surprisingly, although we observed an increase in chromatin density fluctuations, heterochromatic translocation, and gene loci internalization, mesoscale chromatin folding was minimally altered. Indeed, contact scaling, topologically associated domains, chromatin loops, and A/B compartments were minimally perturbed with the joint inhibition of lamin B1 and lamin B2. Further, this effect was similar both within and outside of LAD segments. Likewise, we observed substantial upregulation and downregulation of genes both within and outside of LADs. Phenotypically, changes in expression for genes within LADs were associated with structural disorders and laminopathies (e.g., scoliosis) whereas differentially expressed genes outside LADs were associated with malignancies (e.g., ovarian cancer). Based on these findings, we propose several possible indications: (1) an independent role of B-type lamins in regulation of cellular function independent of mesoscale chromatin regulation and (2) the possibility of a hysteresis in mesoscale chromatin folding induced by association with B-type lamins that is not lost upon joint inhibition. This hysteresis could be due to maintenance of interaction with lamin A/C or direct chromatin interactions with the nuclear envelope.

## RESULTS

### Addition of the mAID tag to B-type lamins does not impact proper localization of lamin A/C

Prior attempts at simultaneous elimination of lamin B1 and lamin B2 have not been successful due to their critical roles in cellular viability and other important cellular processes^13^. To overcome this problem, we utilized the Auxin Inducible Degron system, which allowed simultaneous inhibition of B-type lamins at short timescales^31,32^. In brief, the auxin-dependent degradation pathway found in plants can be introduced into non-plant eukaryotic species to induce rapid depletion of a protein of interest upon exposure to the phytohormone, auxin. This is achieved by fusing a destabilizing domain (degron) to the protein of interest **(Figure 1A-B).** In cells expressing the F-box protein from *Oryza Sativa* (OsTIR1), the addition of auxin results in OsTIR1 forming a functional SCF (Skp1-Cullin-F-box) ubiquitin ligase. Proteins fused with the 7 kDa degron termed mini-AID (mAID) derived from the IAA17 protein of *Arabidopsis thaliana* are rapidly degraded. Importantly, this degradation system is reversible and tunable, allowing for greater control of target protein degradation **(Figure 1D, S1B)**. To achieve this in B-type lamins, we CRISPR-engineered HCT116 colorectal carcinoma epithelial cells and knocked in the mini auxin-inducible degron and mClover at the end of endogenous lamin B1 (LMNB1) and lamin B2 (LMNB2) gene loci **(Figure 1B**) ^33^. For simplicity, we will hereon refer to these engineered cell lines as HCT116^LMNB1-AID^, HCT116^LMNB2-AID^ and HCT116^LMN(B1&B2)-AID^.

**Figure 1:**
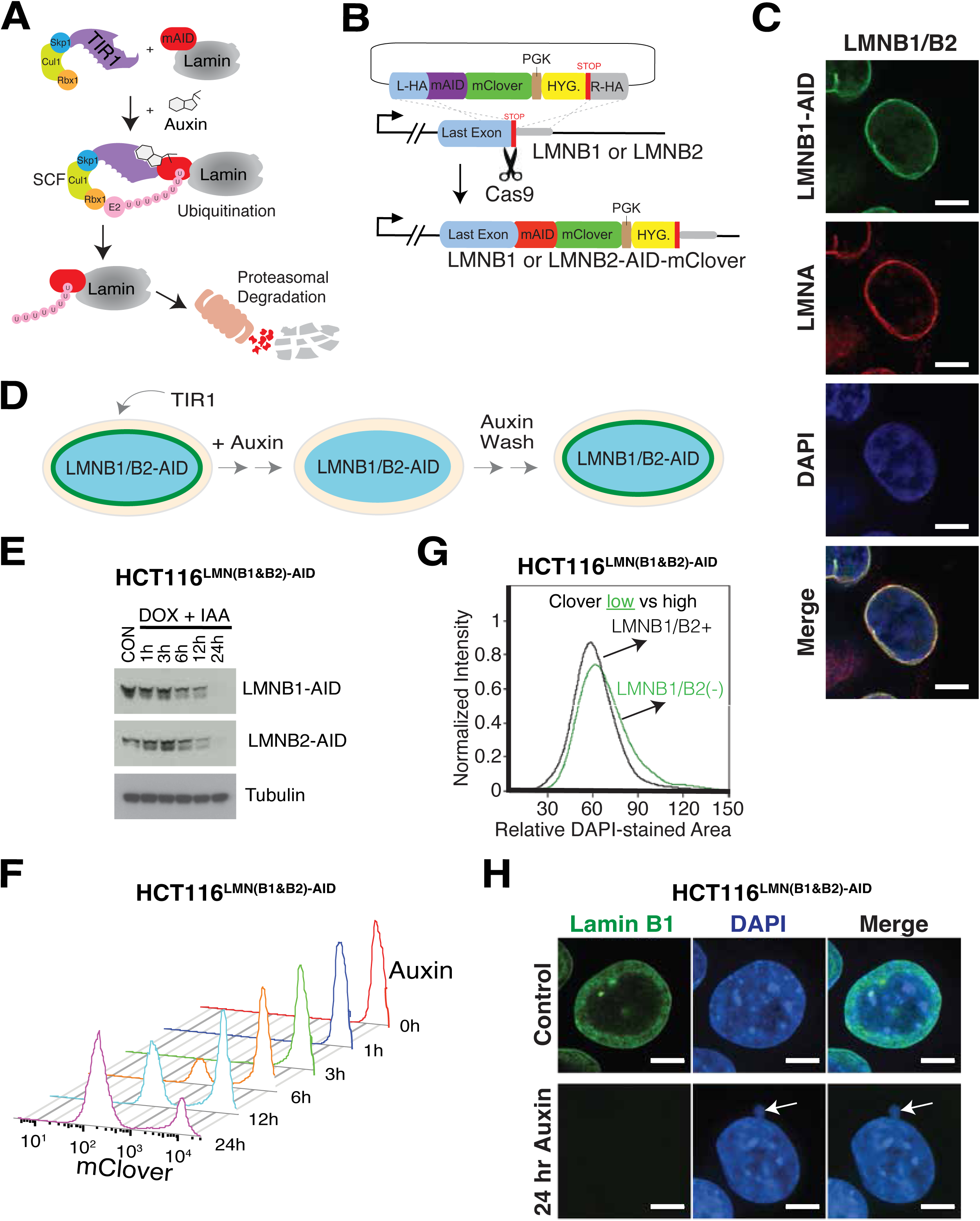
Auxin treatment results in rapid B-type lamin degradation and altered nuclear morphology. A. Auxin treatment promotes the interaction between TIR1 and the degron tag (mAID), which is fused to the target protein. In the presence of auxin, the target protein is rapidly degraded upon proteasomal mediated poly-ubiquitination. B. Each gene of interest was targeted for degradation by co-transfecting progenitor cells with the donor template plasmid with Cas9 and sgRNAs targeting the STOP codon of the sequence. C. The nuclear localization of LMNB1-AID (green) and LMNA (red) in HCT116^LMN(B1&B2)-AID^ cells was examined by immunofluorescence. Cells were stained with DAPI to indicate the nuclear region. Scale bar = 10 μm. N = 3, data was obtained from three independent biological replicates. D. The AID system provides a rapid and reversible method for targeted protein degradation. HCT116^LMN(B1&B2)-AID^ cells express OsTIR1 upon doxycycline addition (left). Auxin treatment results degraded LMNB1 and LMNB2, indicated by loss of mClover (middle). Removal of auxin allows for the re-expression of LMNB1 and LMNB2, indicated by the presence of mClover (right). E. Western blots reveal that AID-tagged LMNB1 and LMNB2 are degraded within 24 hours of auxin treatment, as the target protein expression is no longer detectable in HCT116^LMN(B1&B2)-AID^ cells. F. Single cell-level degradation kinetics demonstrate a loss of B-type lamin proteins. Within 24 hours of auxin treatment, the majority of mClover signal is lost in HCT116^LMN(B1&B2)-AID^ cells. G. Fixed-cell flow cytometry results demonstrate an increase in nuclear area (relative DAPI-stained area) following target protein degradation (green curve), indicated by a low mClover signal, in comparison to untreated cells with high mClover signal (black curve). H. Immunofluorescence shows the presence of nuclear blebbing in B-type lamin-deficient cells. Scale bar = 5 μm. N = 3, data was obtained from three independent biological replicates.

To confirm that addition of the mAID tag degron does not alter the functionality of the tagged lamin proteins, we used immunofluorescence to ensure the continued association of lamin B1/B2 with other proteins that localize to the same compartment as the untagged proteins (i.e., lamin A/C). Genomic localization has previously demonstrated utility as an indicator of functional retention in transcription factor binding ^34,35^. As expected, our results indicate that LMNB1-mAID-mClover, LMNB2-mAID-mClover, and LMNB1/B2-mAID-mClover proteins localize properly to the nuclear lamina in cells **(Figure 1C, S1A)**. We observed a strong overlap between the mClover-signal (hence LMNB1 and/ or LMNB2) and Lamin A/C immunostaining signal. This result suggests that addition of the mAID-mClover tag does not alter proper localization and allows for retention of LMNB1 and LMNB2 protein function.

### Auxin treatment results in rapid and reversible depletion of targeted B-type lamins

We next investigated the kinetics and reversibility of mAID-tagged lamin degradation to confirm the suitability of this approach for reversible inhibition of B-type lamins. A major advantage of the AID system is that conditional depletion of target proteins has less off-target effects than traditional perturbation methods such as RNA interference^31^. This allows examination of resulting changes in chromatin structure and transcription at the most relevant time scales, in which these changes are directly associated with target protein degradation. Further, reversal of target degradation can be achieved by simply removing auxin from cell culture media. We first evaluated the impact of auxin treatment and the reversal process on B-type lamin degradation. To induce the expression of OsTIR1, we added 2 mg/mL doxycycline to cell media 24 hours prior to auxin treatment. Western blot of global protein levels and fixed-cell flow cytometric analysis indicated that 1000 µM auxin treatment resulted in rapid degradation of AID-tagged lamins in HCT116^LMNB1-AID^, HCT116^LMNB2-AID^, and HCT116^LMN(B1&B2)-AID^ cells **(Figures 1E-F, S1C-D)**. Nearly 80% of LMNB1-mAID was degraded within the first 6 hours of auxin treatment. Notably, lamin levels returned to basal levels 3 days after auxin removal. From these results, we confirmed the utility of this mAID system as a means for rapid and reversible degradation of endogenous LMNB1 and LMNB2 proteins to subsequently investigate their role in chromatin folding and cellular function.

### Acute depletion of B-type lamins alters nuclear morphology

As the nuclear envelope is thought to function in the confinement of the genome, we first began by analyzing the effect of B-type lamin inhibition on nuclear morphology. As expected, we observed morphological alterations when either of the lamins is depleted. To quantify these effects, we used ImageStreamX, a flow cytometry based high throughput microscopy imaging platform. At the 8-hour auxin treatment time point for all three cell lines, the population of cells included both mClover-positive (intact lamins) and mClover-negative (degraded lamins) cells. Using this flow cytometry-based analysis of DAPI-stained nuclear area, we found that lamin degradation resulted in enlarged nuclear volume in all three models **(Figure 1G, S2A-B)**. In addition to the change in nuclear volume, confocal sections in HCT116^LMN(B1&B2)-AID^ cells revealed lamin-specific spherical and discrete puncta, which has been previously demonstrated in the nucleus of cultured cells ^36,37^ **(S2C)**. Additionally, we noticed a notable increase in the presence of nuclear deformations, such as nuclear blebbing **(Figure 1H, S1E)**. To evaluate if acute depletion of B-type lamins transformed nuclear morphology due to secondary effects on cell viability, we used Fluorescent-activated cell sorting (FACS) to analyze cell death in HCT116^LMNB1-AID^, HCT116^LMNB2-AID^ and HCT116^LMN(B1&B2)-AID^ cells. Using fluorescently labeled Annexin V (Annexin V^APC^) and DAPI staining to measure overall apoptotic and necrotic cell death,^38^ we confirmed that the acute depletion of both B-type lamins did not induce notable apoptosis or necrosis at 12, 24, or 48 hours of auxin treatment in comparison to the untreated conditions **(S2D)** As such, this indicated that the alterations in nuclear morphology were a direct consequence of B-type lamin removal and that subsequent effects explored were not secondarily due to processes associated with cellular death.

### Acute depletion of both B-type lamins induces minimal changes in mesoscale chromatin structure

Owing to the macroscopic alterations in nuclear morphology observed above as well as prior work showing the role of lamin A/C and LBR in mesoscale chromatin organization, we hypothesized that the depletion of both B-type lamins would produce global alterations in chromatin folding, topologically associated domains, loops, and contact scaling. To test this, we performed *in situ* Hi-C^25^, which can capture structural changes in the kilobase-level DNA-DNA contacts^2^, in the AID cell lines (HCT116^LMNB1-AID^, HCT116^LMNB2-AID^, and HCT116^LMN(B1&B2)-AID^ cells). For each cell type, we generated more than a 1 billion contacts from cells before (∼ 1,354,831,021) and after 24 hours of auxin treatment (∼ 1,210,053,449).

Multiple methods have been proposed for quantifying the mesoscale organization of the genome based on Hi-C data, including (1) analysis of A/B compartment switching^39^, (2) eigenvector decomposition^25^, (3) measurement of contact scaling *(|s|)*^40^, and (4) evaluation of topologically associated domain stability^41–43^. Using Hi-C data, the relationship between chromatin packing behavior and genome connectivity can be conceptualized by the contact probability scaling exponent (*s*). The probability (*P*) of contact between two monomers separated by length (*N*) along a linear chromatin chain follows a power-law scaling relationship: *P ∝ N^-s^*. ^40^ Notably, upon depletion of both lamin B1 and lamin B2, we observed minimal changes in mesoscale chromatin structure **(Figure 2A, S3A-C)**. Depletion of both B-type lamins at 24 hours did not result in major weakening or switching of chromosomal A/B compartments **(Figure 2B)**, markedly alter the frequency or size of topologically associated domains **(Figure 2D, S3D)**, nor change the frequency of contacts as measured by *|s|* **(Figure 2C)**. For example, using TopDom^44^ analysis, a domain-calling algorithm, we found that the mean TAD size slightly decreased (control ∼ 348 kb versus 24 hr. Auxin ∼ 326 kb) while the number of TADs only modestly increased upon 24 hours of auxin treatment (control 7979; 24 hr. Auxin 8491) **(Figure 2D)**. This is in major contrast to the findings observed in LBR depletion where compartment switching, and TADs were markedly transformed in thymocytes lacking LBR compared to WT controls^17^.

**Figure 2:**
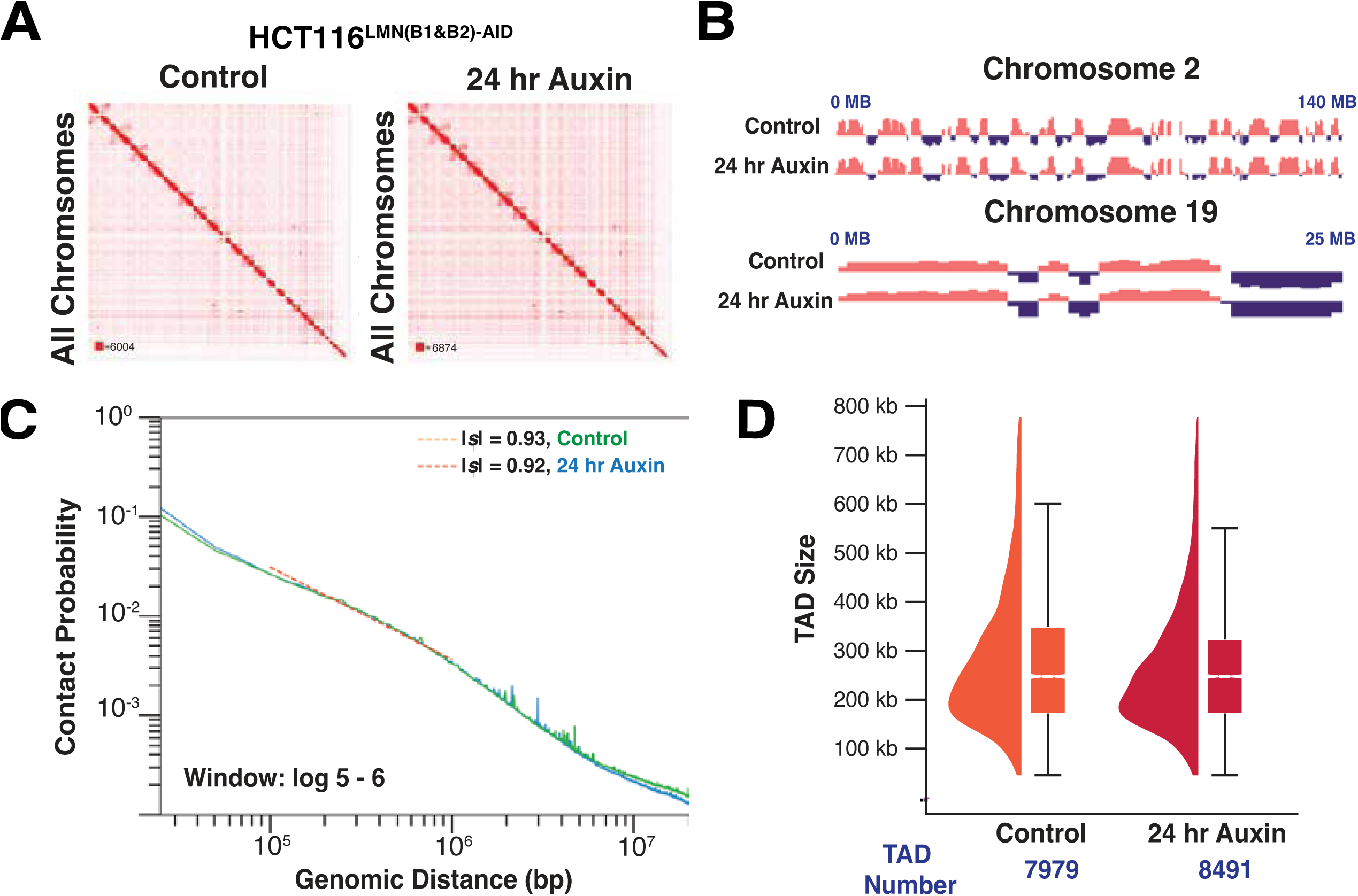
In-situ Hi-C reveals that mesoscale chromatin structure is overall preserved upon B-type lamin degradation. A. Representative normalized Hi-C trans-interaction matrices for all chromosomes in the control and 24-hour auxin treatment conditions are shown for HCT116^LMN(B1&B2)-AID^ cells. B. The eigenvectors for chromosomes 2 and 19 located at the nuclear periphery and interior, respectively, are shown. The A compartment (pink) and B compartment (purple) for each chromosome are indicated. Eigenvectors computed by Juicer are the first eigenvector of the correlation matrix of the binned Hi-C contacts. C. Contact probability scaling for HCT116^LMN(B1&B2)-AID^ cells are shown for the control and 24-hour auxin treatment conditions. Absolute values of *s* are indicated. D. The raindrop plot demonstrates TAD sizes for the control and 24-hour treatment conditions revealed by TopDom. The number of TADs for each condition are indicated below the boxplot.

Next, we hypothesized that one potential explanation of the limited changes in mesoscale chromatin structure upon joint lamin B1 and lamin B2 inhibition could be due to confinement of effects being limited to local alterations around LADs. To perform this segmental analysis, we utilized publicly available DNA adenine methyltransferase identification (DamID) data for Lamin B1 in HCT-116 cells^45,46^. We computationally segmented the genome into contacts that spanned both LADs and non-LADs (across each), across only LADs, or across only non-LADs **(Figure 3A, S3E)**. As prior work has shown that LADs are associated with B-compartments^23,24^, we hypothesized that contact scaling would decay slower within B1 LADs. As expected, we observed a lower |*s*| (calculated between 10^5^-10^6^ bp) within LADs compared to within non-LADs in untreated cells **(Figure 3E)**.

**Figure 3:**
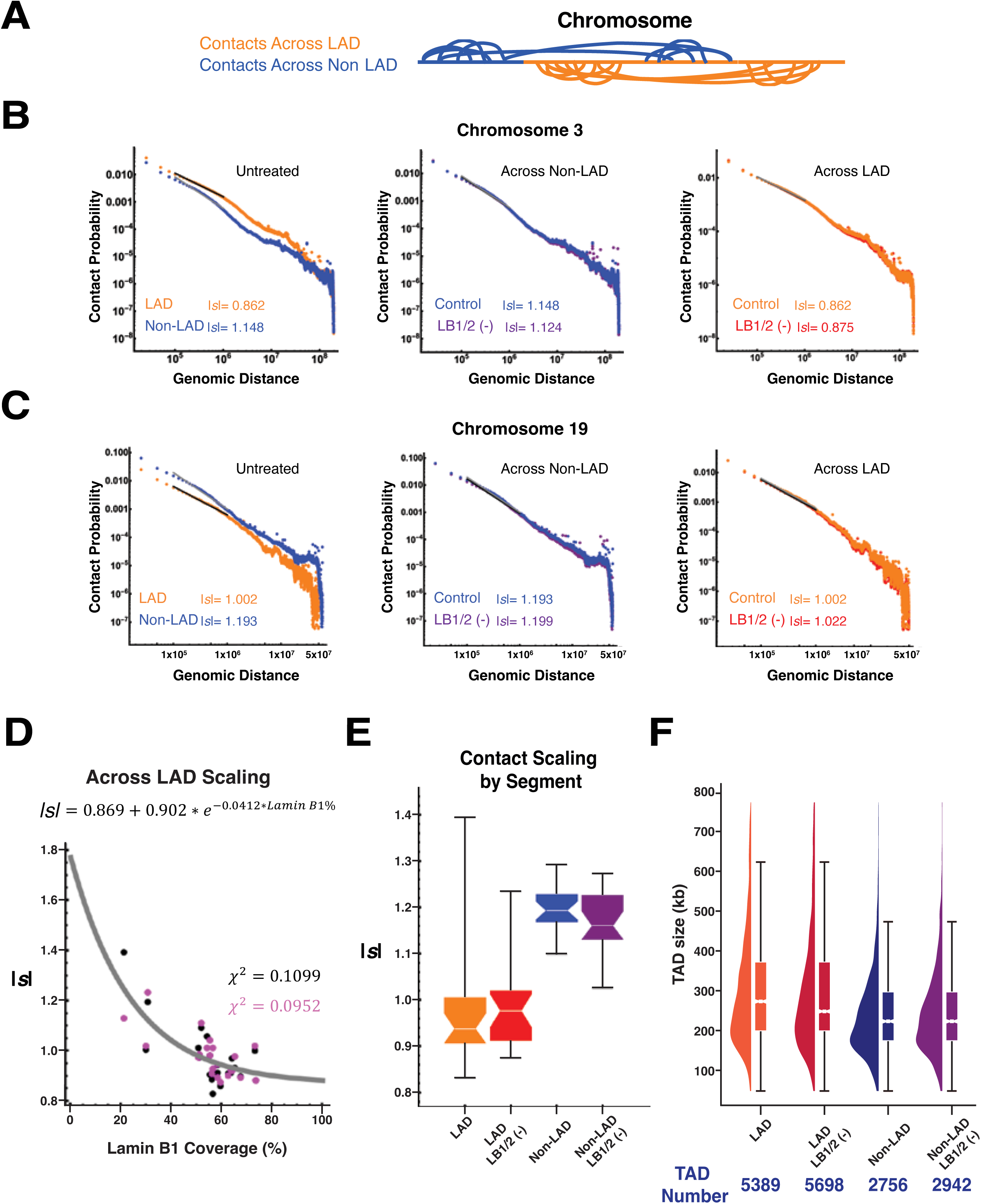
Loss of B-type lamins has a negligible impact on the connectivity of the genome. A. The schematic demonstrates contacts across LAD segments and non-LAD segments for a representative chromosome. B. Contact scaling for chromosome 3 located at the nuclear periphery is shown. The violin plot indicates contact scaling for each chromosome, either across/ within LADs or across/ within non-LADs. Absolute values of *s* are indicated. C. Contact scaling for chromosome 19 located at the nuclear interior is shown. The violin plot indicates contact scaling for each chromosome, either across/ within LADs or across/ within non-LADs. Absolute values of *s* are indicated. D. The scatter plot for the inverse relationship between lamin B1 coverage and |*s|* in LAD segments is shown. The exponential distribution (gray curve) is fitted to the merged control (black) and 24-hour auxin treatment condition (purple) samples. E. The violin plot indicates contact scaling within LADs and non-LADs for both the control (untreated) and 24-hour auxin treatment conditions. F. The rain drop plot demonstrates that TAD sizes do not change in LAD versus non-LAD segments, or in the control versus 24-hour auxin treatment conditions.

When we compared |*s*| for chromosomes associated with the periphery (e.g., chromosome 3) to those associated with the nuclear center (e.g., chromosome 19), we observed a greater difference between |s| in chromosome 3 **(Figures 3B-C)**. Further, analysis of each chromosome demonstrated an inverse relationship between |*s*| within LADs and the percent coverage of the chromosome confined within LADs **(Figure 3D)**. However, despite the large difference in |*s*| at baseline and inverse relationship between |*s*| and LAD coverage, only minor changes in |*s*| were observed upon B-lamin depletion with a larger magnitude of effect observed within the non-LAD portions **(Figure 3E)**. Likewise, as expected, there was no relationship between |*s*| within non-LADs and the chromosomal coverage of lamin B1 **(S3F).** This finding was similarly extended into analysis of TADs. Analysis of TAD size and number within and outside of LADs confirmed that the median size of TADs within LAD regions is higher than TADs outside of LAD regions. However, B-lamin depletion did not differentially impact the size or number of TADs within or outside of LADs **(Figure 3F)**. Overall, these findings indicated that although genomic segments that associate with B-type lamins have a distinct mesoscale structure, the disruption of Lamin B1/B2 surprisingly may not produce phenotypic effects through alterations in higher-order chromatin folding structure.

### Disruption of B-type lamins alters chromatin dynamics but has a minimal effect on higher-order structure in live cells

Although we did not observe major alterations to chromatin folding by *in situ* Hi-C, we hypothesized that disruption of B-type lamins could impart a differential effect on cellular function by changing chromatin dynamics in live cells. To test this hypothesis, we used dual-mode live-cell Partial Wave Spectroscopic (PWS) microscopy to detect both chromatin structural changes and chromatin mobility variations. PWS microscopy provides label-free measurements of nanoscale structural changes with a sensitivity to structures between ∼ 20 and 200 nm in live cells without the use of cytotoxic labels.^47–49^ Briefly, in dual-mode PWS microscopy, variations in nanoscopic-macromolecular structure are measured by analyzing the spectral-dependence in light scattering from the packing of chromatin into higher-order structures, such as chromatin packing domains, while temporal variations in scattering allow measurement of chromatin mobility.^40,50–52^ These domains are characterized by polymeric fractal-like behavior, high chromatin-packing density, and a radial decrease in mass-density from the center to the periphery.^29,53^ As the primary macromolecular assembly within the nucleus is chromatin, this method has been shown to detect changes in chromatin folding comparable to Hi-C and electron microscopy.^40^ Given that higher-order chromatin structure is determined by the polymeric folding across length-scales between 10 – 200 nm and is approximately a power-law (quantified by |*s*| and is observed in multiple polymer models), this distribution in packing corresponds to the distribution of the refractive index quantified by mass scaling (chromatin packing scaling, *D*).^53^ This scaling parameter is calculated for each pixel within the region of interest (i.e., nucleus) within a given coherence volume, which is determined by the depth of field longitudinally and the axial plane for each pixel. Therefore, spatial variations of macromolecular density and motion that can occur upon perturbation, such as degradation of B-type lamins, can be measured using PWS.

First, we confirmed with PWS that OsTIR1 expression (upon doxycycline treatment) does not alter *D* **(S4E)**. As |*s*| and *D* are inversely related based on prior experimental studies and polymer modeling^40,52^, we hypothesized that that nuclear average *D* would increase slightly after inhibition of lamin B1 and lamin B2 due to the small decrease in |*s*| observed within non-LAD portions of the genome.^40^ Upon the addition of auxin to HCT116^LMN(B1&B2)-AID^ cells, the average *D* increased by a minimal amount of 0.8% but statistically significant amount across the nucleus (0.021 ± 0.002 (SEM), *n* = 1822, *p-value* < 0.0001) **(Figure 4A-B)** with larger but still modest changes observed within HCT116^LMNB1-AID^ and HCT116^LMNB2-AID^ **(S4A-B,** *n =* 482 & 401, respectively). We verified a strong correlation between the change in *D* and mClover signal with a 24-hour auxin treatment time course **(S4E)**. Further, the magnitude of this effect was substantially smaller than that observed in other conditions, such as in transcription inhibition by actinomycin D in BJ fibroblast cells (7% decrease).^40^ Overall, the small increase in *D* and decrease in |*s*| suggested that the regulatory role of nuclear B-type lamins was independent of mesoscale chromatin folding in both live and fixed cells.

**Figure 4:**
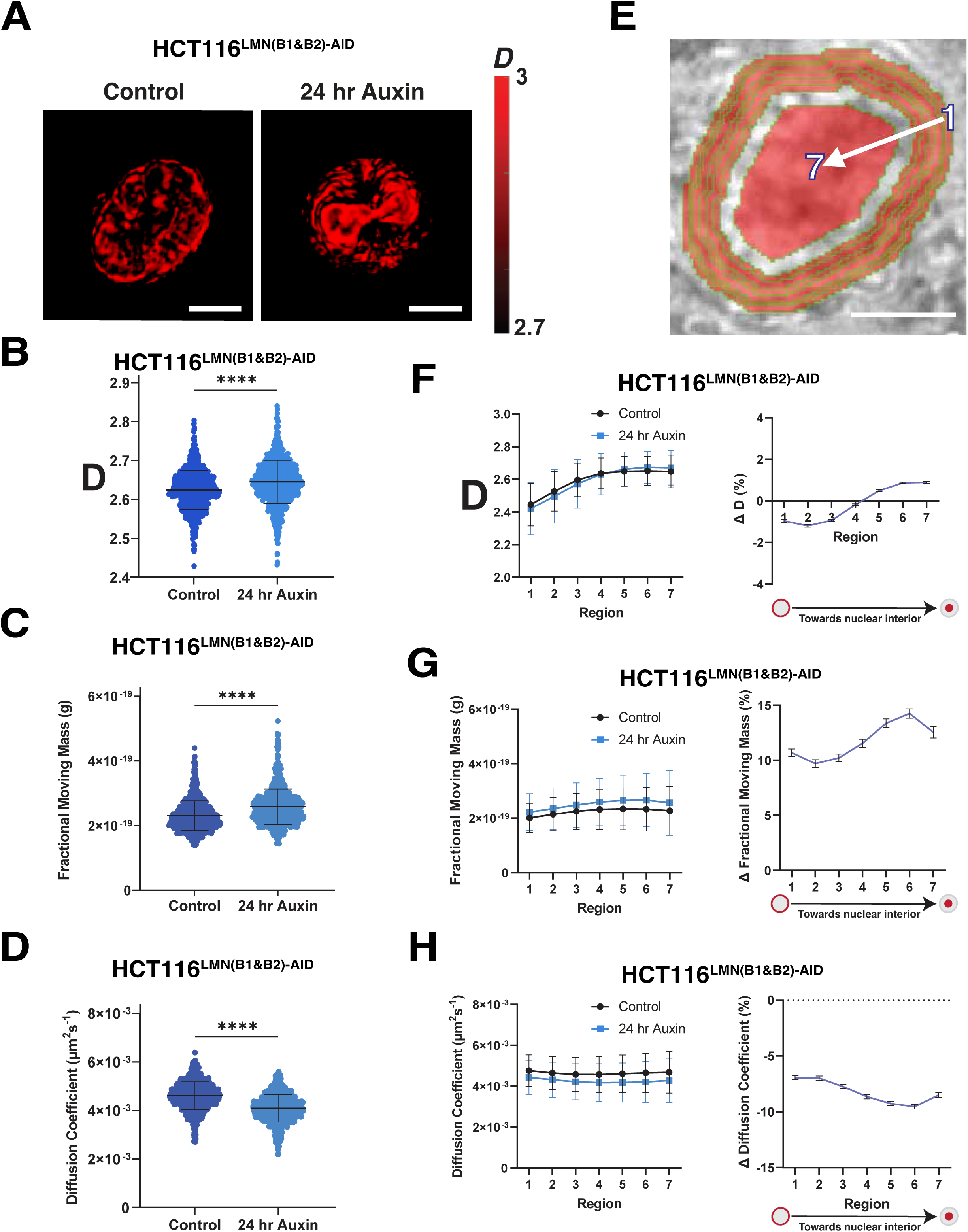
Dual-PWS microscopy reveals differential higher-order chromatin structure and dynamics upon loss of B-type lamins. A. Representative images of HCT116^LMN(B1&B2)-AID^ cells obtained from PWS are shown for both the control and 24-hour auxin treatment conditions. Scale bar = 5 μm. N = 3, data was obtained from three independent biological replicates. B. Violin plots demonstrate that chromatin packing scaling (D) is significantly increased in HCT116^LMN(B1&B2)-AID^ cells. C. Violin plots demonstrate that fractional moving mass is significantly increased in HCT116^LMN(B1&B2)-AID^ cells upon 24 hours of auxin treatment. D. Violin plots demonstrate that the diffusion coefficient is significantly decreased in HCT116^LMN(B1&B2)-AID^ cells upon 24 hours of auxin treatment. (B-D). The truncated violin plots extend from the minimum to the maximum value. The line in the middle of each plot is the median value of the distribution, and the lines above and below are the third and first quartiles, respectively. Data was obtained from three technical replicates for each condition. ****P<0.0001. Welch’s correction was applied. E. The schematic indicates that region 1 is the nuclear periphery while region 7 is the nuclear interior. Scale bar = 5 μm. F. Regional PWS measurements of *D* HCT116^LMN(B1&B2)-AID^ cells are shown, along with the change in *D* within each region. G. Regional PWS measurements of fractional moving mass for HCT116^LMN(B1&B2)-AID^ cells are shown, along with the change in fractional moving mass within each region. H. Regional PWS measurements of the diffusion coefficient for HCT116^LMN(B1&B2)-AID^ cells are shown, along with the change in the diffusion coefficient within each region.

Utilizing the temporal capabilities of dual-PWS, we then evaluated the influence of B-type lamin degradation of chromatin dynamics^50,51^. Although dual-PWS lacks molecular specificity, variations in temporal interference from macromolecular movement allow quantification of the fractional moving mass (the magnitude of chromatin evolving in time) and the diffusion coefficient without labels. Fractional moving mass, calculated from the product of the mass of the moving macromolecular cluster and the volume fraction of moving mass, quantifies the physical and dynamic properties chromatin, as not all motion within the cell is specifically diffusive.^50^ Unexpectedly, although minor changes in higher-order chromatin structure were observed upon lamin B1/B2 depletion, there were much larger changes in the fractional moving mass on treatment with auxin in HCT116^LMN(B1&B2)-AID^ cells (18.4%) (2.780 ± 1.275e-21 (SEM), *n* = 1822, *p-value* < 0.0001) **(Figure 4C).** Similar changes were observed in HCT116^LMNB1-AID^ and HCT116^LMNB2-AID^ **(S4C).** These results suggest that upon lamin B1 and lamin B2 degradation, there is an increase in molecular motion occurring universally within the cell across a large range of dynamic processes.^50^ Indeed, we further observed that inhibition of lamin B1 and lamin B2 resulted in a decrease in the ensemble diffusion coefficient of 11.4% compared to the untreated control (0.000 ± 2.664e-5 (SEM), *n* = 1822, *p-value* < 0.0001) **(Figure 4D)**, with similar findings in HCT116^LMNB1-AID^ and HCT116^LMNB2-AID^ **(S4D).** Taken together, our temporal analysis indicates that B-type lamin degradation results in overall greater magnitude of chromatin turnover with a slower rate of motion.

### Lamins B1 and B2 mechanistically determine chromatin mobility and heterochromatin localization throughout the nucleus

Based on our observations above on the slight differential effect in *|s|* in LAD vs non-LADs and the alterations in chromatin mobility, we investigated if the mechanistic role of B-type lamins is in the stabilization of chromatin cores to the nuclear periphery. To distinguish between whole-nuclei and regional chromatin structure, we segmented the nucleus into regions of equal size spanning the nuclear edge and the center of the nuclear interior **(Figure 4E)**. This spatial analysis involved segmenting the nucleus into six non-overlapping ribbons to evaluate variations in the location of chromatin packing domains. Structurally, *D* near the nuclear periphery was lower than that of the nuclear interior, with minimal differences observed between the periphery and the interior during lamin B1 lamin B2 inhibition **(Figure 4F)**, with similar trends observed in Lamin B1 and B2 inhibition separately **(S4F)**. Further, *D* was strongly correlated with the distance from the periphery (Pearson correlation coefficient = 0.95). This differential spatial response of the change in *D* could potentially be explained by the movement of chromatin between the periphery and nuclear interior. Unexpectedly, although fractional moving mass was increased and diffusion was decreased throughout the nucleus, the differences between the nuclear periphery and the center in HCT116^LMN(B1&B2)-AID^ cells was minimal **(Figures 4G-H)**, with comparable results observed in HCT116^LMNB1-AID^ and HCT116^LMNB2-AID^ **(S4G-H)**.

Overall, these findings suggested that the mechanistic role of lamin B1 and lamin B2 was in confinement of chromatin mobility throughout the cell nucleus and suggested that when lamin B1 and lamin B2 are removed, chromatin moves away from the nuclear periphery with rippling effects into the nuclear interior regions. We reasoned that if LMNB1 and LMNB2 depletion leads to the detachment of chromatin from the nuclear periphery, as indicated by our dynamic PWS results, this shift would be accompanied by heterochromatin redistribution and peripheral decompaction. Using a combination of immunofluorescence and western blot analysis, we confirmed an internal shift and global decrease in heterochromatin (H3K9me2/3 and H3K27me3) and an increase in euchromatin (H3K27ac) upon 24 hours of Auxin treatment **(Figure 5A-B, S5A-C)**. To estimate changes in chromatin compaction upon the removal of B-type lamins, we used spinning disk confocal microscopy to measure the coefficient of variation in DAPI-stained HCT116^LMN(B1&B2)-AID^ cells. Previously used to assess the degree of heterogeneity of DNA signal across the nucleus^54^, the coefficient of variation is calculated as the standard deviation of the DAPI intensity values divided by the mean value of nuclear pixel intensity for each nucleus. Our results demonstrated that upon auxin treatment, chromatin compaction slightly decreased in HCT116^LMNB1-AID^ (1.11%), HCT116^LMNB2-AID^ (2.31%), and HCT116^LMN(B1&B2)-AID^ (2.86%) cells **(Figure 5C, S5E-F)**. Therefore, the removal of B-type lamins results in destabilization of heterochromatin localization. Taken together, these results indicate that although mesoscale organization is maintained, the shift in heterochromatin at the nuclear periphery induces global nuclear changes due to alterations in chromatin mobility.

**Figure 5:**
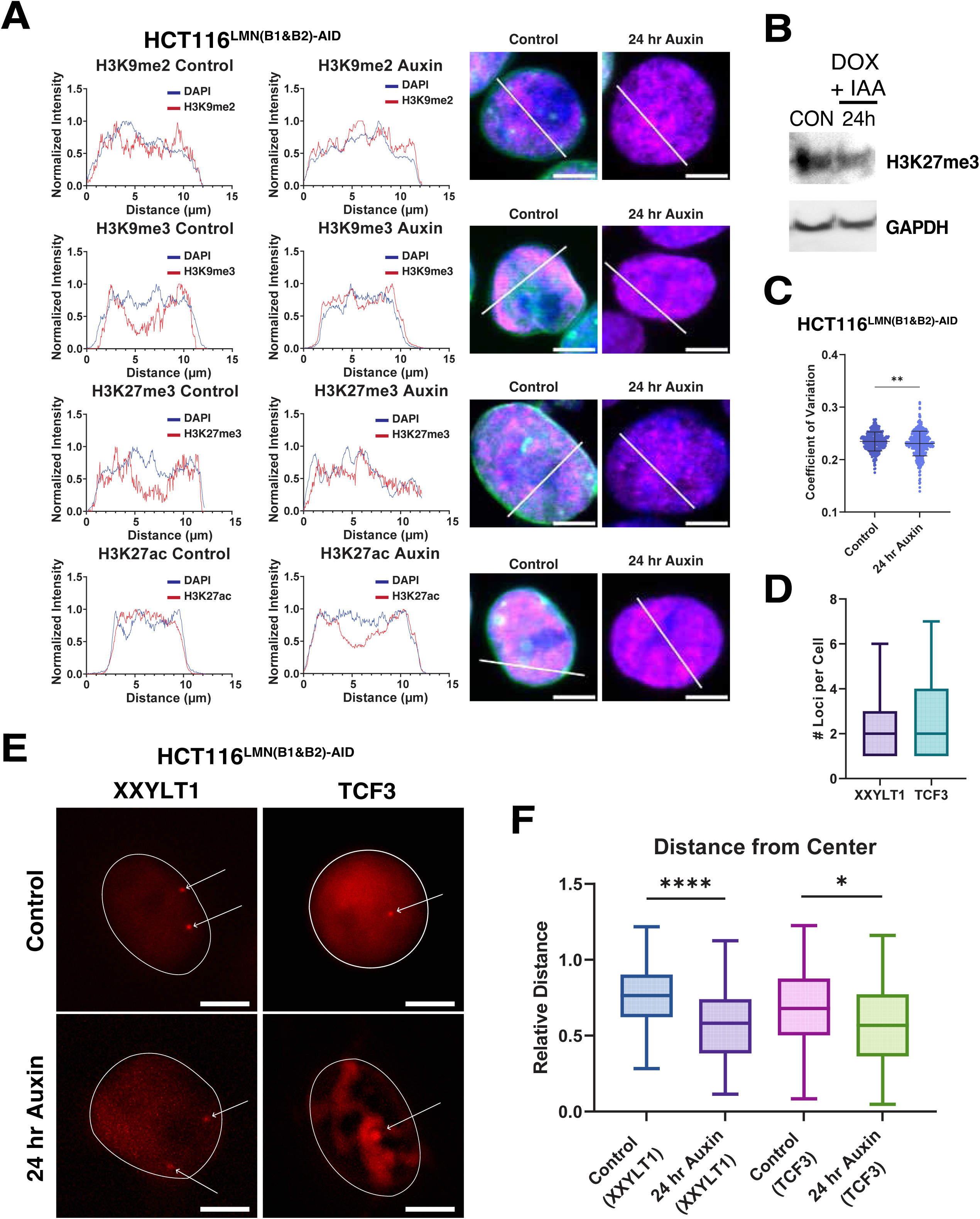
Positioning of heterochromatic markers, chromatin compaction, and chromosome positioning are significantly altered upon auxin treatment. A. Line plots for the distribution of heterochromatic and euchromatic marks are shown (red) in comparison to DAPI signals (blue) for the representative nuclei from spinning disk confocal microscopy (Lamin B1/B2 in green, histone marks in magenta, DAPI in blue). B. Western blots reveal that H3K27me3 expression decreases by ∼25% within 24 hours of auxin treatment. Within this time frame, the target protein expression is no longer detectable in HCT116^LMN(B1&B2)-AID^ cells. C. The coefficient of variation plot for HCT116^LMN(B1&B2)-AID^ cells demonstrates chromatin decompaction upon 24 hours of auxin treatment. The truncated violin plots extend from the minimum to the maximum value. The line in the middle of each plot is the median value of the distribution, and the lines above and below are the third and first quartiles, respectively. Data was obtained from three technical replicates for each condition (N=3, Control (*n* = 798), 24-hour auxin (*n* = 845)). ****P<0.0001. Welch’s correction was applied. D. Box plot showing the number of foci per cell for XXYLT1 and TCF3 loci; *n*□=□42 cells (XXYLT1) and 25 cells (TCF3). The line within each box represents the mean; the outer edges of the box are the 25th and 75th percentiles and the whiskers extend to the minimum and maximum values. E. Representative images of gene loci XXYLT1 and TCF3 in live cells. B-type lamin depletion alters the subnuclear localization preference of genomic loci, mostly at the nuclear periphery. Scale bar = 5 μm. F. Box plot showing distances of XXYLT1 and TCF3 loci to the center *n*□=□53 cells (XXYLT1) and *n=* 28 cells (TCF3). The line within each box represents the mean; the outer edges of the box are the 25th and 75th percentiles and the whiskers extend to the minimum and maximum values. ****P<0.0001.

### Auxin treatment results in lamina-dependent changes in 3D spatial localization and dynamic chromatin reorganization

Owing to the differential effect of LB1/B2 depletion on chromatin mobility and spatial distributions observed in HCT116^LMN(B1&B2)-AID^, we next tested if re-localization of heterochromatin was paired with gene-re-localization from the nuclear periphery. Specifically, although Hi-C demonstrated muted changes in |*s*|, compartments, and TADs, it remained possible that heterochromatin cores were moving independent of the relocation of genes in 3D space. To test this, we utilized CRISPR-Sirius, a DNA imaging system for imaging of chromosome-specific loci with a high signal-to-noise ratio in live cells ^55^ to measure the spatial distribution of genes in live cells upon lamin B1 and lamin B2 depletion. Since depletion of both lamins B1 and B2 resulted in nuclear wide changes in mobility and heterochromatin localization while mesoscale structures were maintained, we hypothesized that gene loci would similarly relocate in response in HCT116^LMN(B1&B2)-AID^ upon auxin treatment.

To assess the effect of B-type lamin degradation on chromosome-specific loci in live HCT116^LMN(B1&B2)-AID^ cells, we labelled genomic regions on chromosomes 3 and 19 using CRISPR-Sirius. Chromosomes 3 and 19 were chosen to represent chromosomes that are localized relatively near the nuclear periphery and the nuclear interior, respectively.^55–60^ Specifically, we chose one intronic gene region on Chromosome 3 (*XXYLT1*) containing ∼ 333 tandem repeats, and one intronic gene region on Chromosome 19 (*TCF3*) containing ∼ 36 tandem repeats as a reference to the nuclear interior. To measure the spatial distance and dynamics of the *XXYLT1* gene loci, we labeled them with CRISPR-Sirius-8XMS2 and the MS2 capsid protein MCP-HaloTag, which contains a nuclear localization sequence.^55^ We visualized 2-3 pairs of loci in individual cells **(Figure 5D-E)** and quantified the average distance between the nuclear periphery and either the *XXYLT1* or *TCF3* gene loci, normalized to the nuclear radius **(Figure 5F)**. Our results demonstrated that the average distance between the nuclear periphery and *XXYLT1* significantly increased upon lamin degradation, indicating that the targeted loci moved towards the nuclear interior, as expected. In conjunction with relocation of heterochromatin and nuclear wide changes in chromatin mobility, the average distance between the nuclear periphery and TCF3 also increased, although to a less significant degree. These results indicate that upon B-type lamin degradation, chromatin reorganizes throughout the nuclear area, especially at the nuclear periphery.

### Acute B-type lamin depletion alters gene expression within both LADs and Non-LADs

Although the changes in mesoscale chromatin structure were muted, we hypothesized that alterations in chromatin localization and dynamics could still result in profound changes in gene expression. To test this, we utilized RNA-Sequencing to measure gene expression changes between untreated cells to 12-hour auxin-treated cells, 48-hour auxin-treated cells, and cells that were auxin-treated for 48 hours followed by a 6-day wash off **(S6A)**. Given the global changes in nuclear organization observed, we hypothesized that depletion of B-type lamins would result in differential gene expression within both LAD and non-LAD territories. We anticipated that genes within LADs would experience upregulation once freed from the heterochromatic nuclear periphery. Likewise, as mesoscale structure was preserved, we anticipated that there would be limited association between gene expression changes and TAD structure. To identify broad transcriptomic phenotypes following LMNB1/B2 degradation, we analyzed differentially expressed genes (DEGs) at each timepoint **(S6B)**. B-type lamin depletion led to widespread differential expression: 887 genes were upregulated, and 830 genes were downregulated after 48 hours of auxin treatment, when the effect of lamin depletion was most penetrant (adjusted P value < 0.01 and absolute log fold change > 1). Further, restoration of B-type lamins after growing HCT116^LMN(B1&B2)-AID^ cells in auxin-free media for 6 days after 48 hours of auxin treatment returned cells to basal levels of gene expression. To summarize, these findings indicate that B-type lamins have a distinct mechanistic role in transcriptional regulation that is independent of mechanisms responsible for maintaining higher-order chromatin folding.

Based on prior work, we anticipated that many of the differentially expressed genes after 48 hours of B-lamin depletion would be found within LADs. Using the same publicly available Lamin B1 DamID data from HCT116 cells, we again segmented the genome into regions within LADs and regions outside LADs^45^. After 48 hours of lamin depletion, we identified nearly 22% of enriched genes within LADs (adjusted P value < 0.01 and absolute log fold change > 1) **(Figure 6A)**. Anomalously, lamin B1 and lamin B2 depletion predominantly generated the highest number DEGs outside of LAD boundaries for all three timepoints, with the greatest number at 48 hours of auxin treatment (374 genes within LADs, 1331 genes outside of LADs at 48 hours) **(S6C-D)**. Unexpectedly, in both LAD and Non-LAD regions, there was no clear trend towards upregulation or downregulation in gene expression. Further, the log fold change of DEGs outside of LADs was larger in magnitude than within LADs, indicating that they were disproportionally influenced by loss of B-type lamins **(S6E)**.

**Figure 6:**
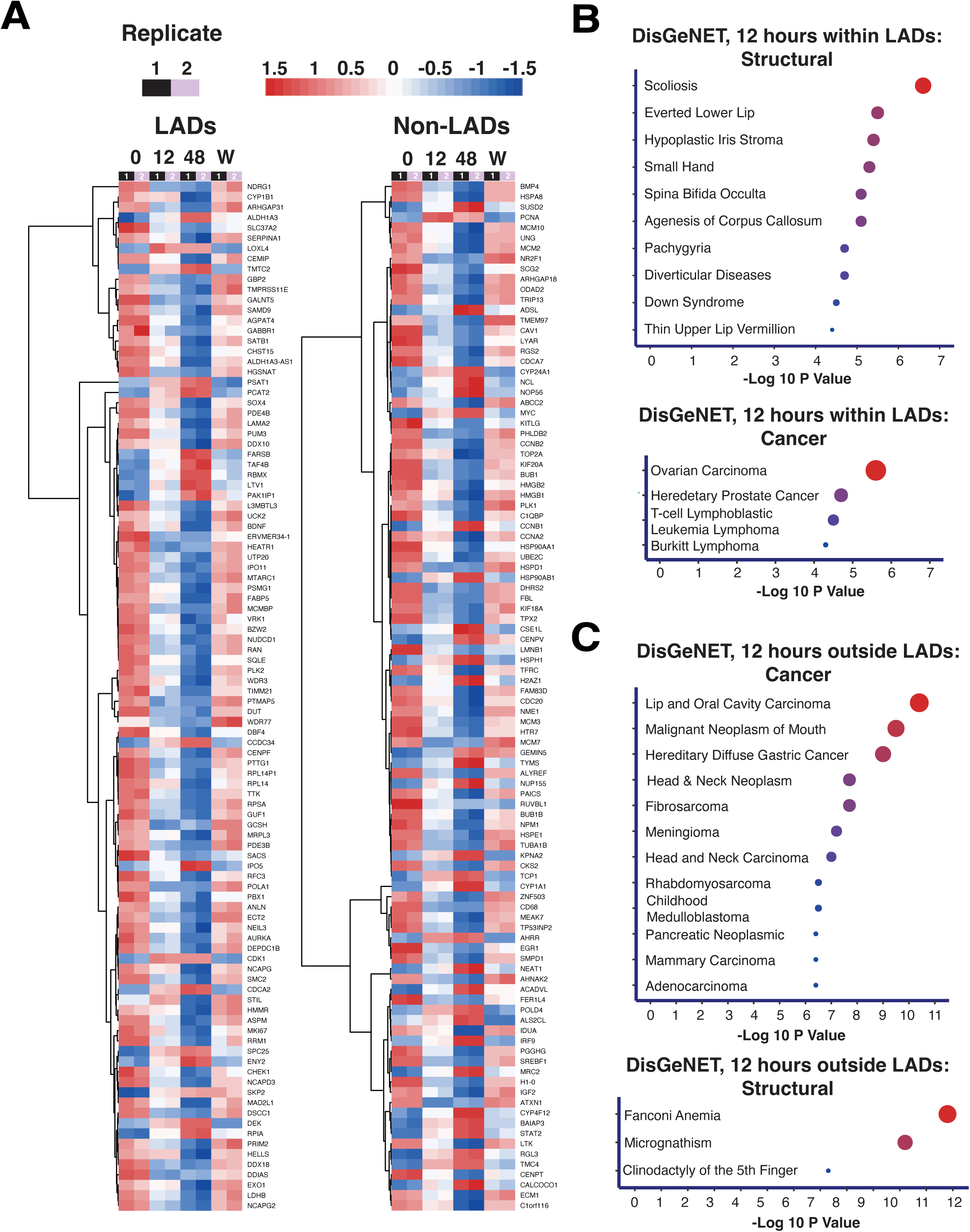
Loss of both B-type lamins perturbs the expression of genes positioned inside and outside LADs. A. Heatmaps showing the top 100 DEGs within versus outside of LAD boundaries upon 48 hours of auxin treatment defined by publicly available DamID data (adjusted P value < 0.01 and absolute log fold change > 1). B. DisGeNET results indicate terms upon 12 hours of auxin treatment within LADs related to structural changes and cancer. C. DisGeNET results indicate terms upon 12 hours of auxin treatment outside of LADs related to cancer and structural changes.

**Figure 8:**
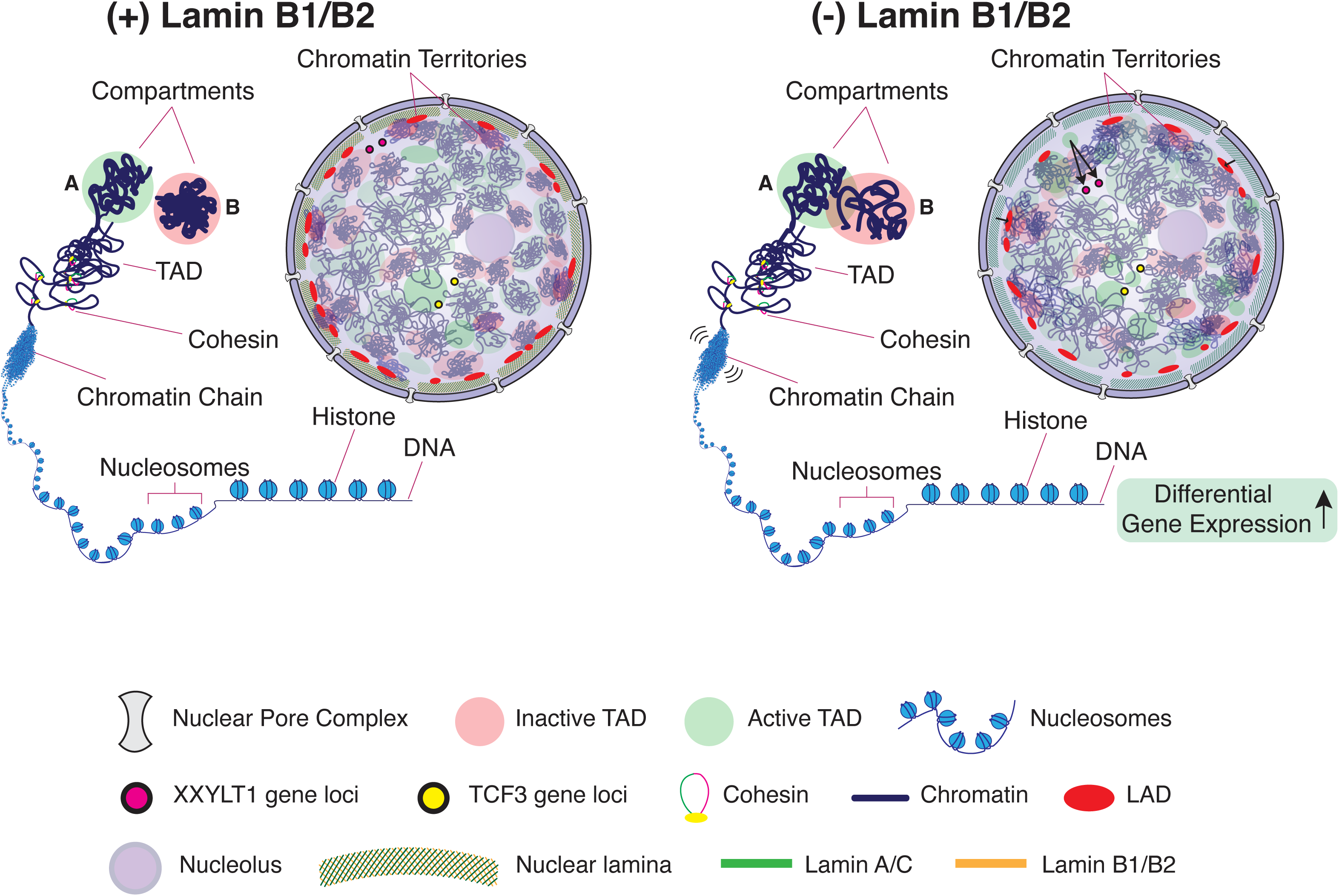
Proposed model of the effect of Lamin B1 and B2 degradation on chromatin organization. Schematic depicting how inhibition of B-type lamins leads to a shift in higher-order chromatin structure and dynamics. Loss of lamins B1 and B2 promotes chromatin blending, chromatin chain fluctuations, and displacement of both heterochromatin-associated domains and chromosome-specific loci, especially at the nuclear periphery. Gene expression is dramatically altered as a function of heterochromatin reorganization upon the loss of structural constraints at the nuclear periphery.

With respect to the properties of DEGs within LADs and Non-LADs, we performed a gene ontology analysis using Metascape and the Disease Gene Network Database. Notably, DEGs within LADs were enriched for a handful of cancers and laminopathies such as scoliosis **(Figures 6B-C)**. Together, these data suggest that LADs maintain transcriptionally quiet states for many genes and upon depletion of B-type lamins, unevenly upregulate genes leading to altered transcriptional states that contribute to laminopathies and cancers. We also analyzed the transcriptional divergence of genes within and outside of LADs. Transcriptional divergence is a measure of the change in expression between minimally expressed transcripts and highly expressed transcripts. Using the Gini equation, a common metric borrowed from mainstream economics for measuring equality across a population, we calculated the Gini coefficients for each timepoint to quantify transcriptional divergence^61^. Upon 48 hours of auxin treatment, the Gini coefficient for all genes decreased modestly from ∼ 0.87 to 0.85. We also found that transcriptional divergence was generally higher within LADs versus non-LADs **(S6F)** and that genes within LADs were more heterogeneously expressed upon B-type lamin depletion **(S6E)**.^62^ Given that DEGs within LADs were associated with laminopathies and structural disorders while DEGs in non-LADs were primarily associated with malignancy, these results indicate distinct mechanistic consequences of gene positioning.

## DISCUSSION

In this work, we generated clonal cell lines where endogenous copies of LMNB1 and LMNB2 were fused with mAID to achieve rapid and reversible degradation of B-type lamins in the cell through auxin exposure. Using this approach, we investigated the role of B-type lamins on chromatin organization, dynamics, cell function, and gene expression. Our results demonstrate that auxin treatment results in an enlarged nuclear area and can effectively deplete B-type lamins within 24 hours without resulting in apoptosis or necrosis. The consequence of lamin B1 and lamin B2 depletion had surprising effects on nuclear structure and gene expression. Critically, our findings indicate that the functional consequences of B-type lamin degradation do not depend on changes to the mesoscale chromatin structure (i.e., TADs, contact scaling, or compartments), but rather on alterations in chromatin dynamics and the spatial distribution of the genome. However, lamins regulate spatial chromatin organization as we observe that lamin depletion leads to the movement of gene-specific loci towards the nuclear interior and increased density fluctuations in the chromatin polymer.

In recent Hi-C studies comparing conventional and inverted nuclei that form from LBR depletion or Lamin A/C depletion, relatively few changes in compartment and chromatin connectivity are observed^17^. While B-type lamin depletion leads to shifts in gene expression within and outside of LADs, our results demonstrate that these changes are not from compartment switching, TAD depletion, or changes in chromatin packing efficiency. Instead, lamin B1 and lamin B2 seem to exert their effect on chromatin structure independent of the regulatory mechanisms confining the genome at the mesoscale. Although B-type lamin depletion led to increased nuclear volume, the overall maintenance of loops, TADs, and compartments indicates that genome topology is robust to perturbation of chromatin elasticity.^43^ These modest effects on higher order chromatin structure could be due to multiple mechanisms, including (1) redundancy of LMNA proteins after depleting LMNB1 and LMNB2, (2) a distinct role for LBR in chromatin organization, or (3) the evolution of hysteresis in chromatin marks due to association with B-type lamin domains that is not lost on their disruption.

Our findings highlight the need for development of methods for the investigation of chromatin dynamics as well as evolution in chromatin modeling to better understand how the temporal evolution of chromatin can alter gene transcription without alterations in mesoscale structure. While Hi-C methods enable the reconstruction of 3D chromosome and genome structures, one of the key limitations of this technique is that that long-range frequencies can be noisy and unreliable, often depending on experimental factors such as the length of restriction fragments, GC content, and read depth.^63^ Further, this is a fixed-cell technique and thus does not allow for millisecond resolution of chromatin structure and dynamics. Consequently, Hi-C and derivative techniques are incapable of detecting the complexities of dynamic chromatin reorganization alone. Thus, incorporating multi-modal imaging techniques in addition to Hi-C can provide a better method for characterizing nanoscale structural alterations in chromatin packing in real time.

Future work pairing super-resolution microscopy in combination with PWS microscopy and high-resolution electron microscopy, involving labeling for markers specific to heterochromatin, could help elucidate the nanoscopic changes in chromatin structure during lamin B disruption.^30,40^ Such an approach can also aid in providing further understanding of the mechanisms that maintain the phenotypes involving degraded lamins or inverted nuclei. Despite these limitations, our findings provide a deep and surprising role of B-type lamins in cell type-specific 4D genome organization and transcription. In this work, increased fluctuations in density appear to be associated with the translocation of genes spatially and translocation of heterochromatin markers. Theoretical work regarding fluctuations in density has been previously shown to independently regulate gene expression.^64^ Upon lamin degradation, heterochromatic marks slightly shifted from the nuclear periphery towards the interior, indicating greater variability in accessible chromatin loci.

The functional consequence of the transformation in chromatin organization on gene expression patterns occurs both for genes within and outside of LADs upon depletion of LMNB1 and LMNB2. As such, our work provides further evidence that LADs function not solely through the effect on transcriptional repression but are involved in the maintenance of transcriptionally active genomic sites. Contradictory findings from previous studies may be due to several factors. First, some LADs are cell-type specific, whereas others are conserved (i.e., constitutive LADs).^23,24,62^ This could impact the resulting changes seen in genome-nuclear lamina interactions due to changes in gene expression (lamin perturbation). Further investigation in other cell types could provide a method to better understand how lamins maintain cell-type specific structural organization in normal and disease development. For example, A-type lamins and perinuclear lamin B2 have been previously shown to be involved in age-related neurodegeneration.^65^ Our results therefore warrant additional studies using neurons or iPSCs utilizing similar inhibitory mechanisms presented here.

## CONCLUSION

Our investigation focuses on understanding the mechanisms by which the rapid inhibition of both B-type lamins simultaneously would alter genome organization, gene expression, and cell function. Our results in mammalian cells indicate a direct link between transcription changes mediated by B-type lamins near LADs that is not due to higher-order chromatin structure transformation. Transcriptionally, expression in both LAD and non-LAD territories are transformed, with distinct disease associations from the expression patterns depending on the genomic territory (LADs associated with structural disorders and non-LADs with malignancies). Further, our results demonstrate that upon B-type lamin degradation, chromatin conformation and chromatin connectivity show subtle changes. However, the greatest alterations in chromatin conformation and connectivity occur closest to the nuclear periphery in comparison to the nuclear interior. Consequently, these findings demonstrate a distinct mechanistic role of B-type lamins in chromatin organization and cellular function. Our results therefore warrant further investigation of the dynamic relationship between the nuclear lamina and chromatin organization.

## MATERIALS AND METHODS

For more details, see Key Resources Table.

### HEK293T Cell Culture

HEK293T cells (ATCC, #CRL-1573) were grown in Dulbecco’s Modified Eagle’s Medium (DMEM) supplemented with 10% FBS (#16000-044, Thermo Fisher Scientific, Waltham, MA) and penicillin-streptomycin (100 μg/ml; #15140-122, Thermo Fisher Scientific, Waltham, MA). All cells were cultured under recommended conditions at 37°C and 5% CO_2_. All cells in this study were maintained between passage 5 and 20. Cells were allowed at least 24□h to re-adhere and recover from trypsin-induced detachment. All cells were tested for mycoplasma contamination (ATCC, #30-1012K) before starting experiments, and they have given negative results.

### HCT116 Cell Culture

HCT116 cells (ATCC, #CCL-247) were grown in McCoy’s 5A Modified Medium (#16600-082, Thermo Fisher Scientific, Waltham, MA) supplemented with 10% FBS (#16000-044, Thermo Fisher Scientific, Waltham, MA) and penicillin-streptomycin (100 μg/ml; #15140-122, Thermo Fisher Scientific, Waltham, MA). All cells were cultured under recommended conditions at 37°C and 5% CO_2_. All cells in this study were maintained between passage 5 and 20. Cells were allowed at least 24□h to re-adhere and recover from trypsin-induced detachment. All imaging was performed when the surface confluence of the dish was between 40–70%. All cells were tested for mycoplasma contamination (ATCC, #30-1012K) before starting perturbation experiments, and they have given negative results.

## METHOD DETAILS

### Transfection and Colony Isolation for Creating AID cell lines

Stable transfection was achieved as previously described ^33^. Briefly, we generated conditional human HCT116 mutants by homology-directed repair (HDR)-mediated gene tagging using CRISPR-Cas9. Around 60-70% confluent cells expressing OsTIR1 were plated at 3×10^5^ cells in a 6-well plate and cultured for 24 hours at 37°C. On the next day of transfection, 4 mL of 200 ng/mL CRISPR plasmid, 3 mL of 200 ng/mL donor plasmid, 90 mL of Opti-MEM I Reduced Serum Medium (Gibco, #31985070), and 8 mL of FuGENE 6 Transfection Reagent (Promega, #E2691) were mixed and incubated at room temperature for 15 minutes before being applied to the cells. For antibiotic selection and colony formation, Hygromycin B Gold (Gibco, #10687010), 100 mg/mL was used. For colony isolation, single colonies were picked under a stereo microscope and transferred to a 96-well plate containing 10 mL of trypsin/ EDTA and neutralized with 200 mL of media. After 2 weeks of culture, the cells were transferred to a 24-well plate. After a few days of culture, genomic DNA was isolated. Briefly, the cells were harvested, lysed with SDS buffer (100 mM NaCl, 50 mM Tris-Cl pH 8.1, 5mM EDTA, and 1% wt/vol SDS) and treated with proteinase k (New England Biolabs, #P8102) (0.6 mg/mL) at 55□°C for 2 hours. Then, the lysis solutions were treated with PCl (phenol/ chloroform/ isoamyl alcohol) and used in EtOH precipitation. Genomic DNA pellets were washed with 70% EtOH and resuspended in RNase-containing water. The PCR reaction was set up using 0.5 U of Taq DNA Polymerase (G Biosciences, #786-447) (1x PCR Buffer), 0.5 mM primers, and 1 mL of genomic DNA from the HCT116 CMV-OsTIR1 parental cells to a 20 mL total volume reaction mixture. PCR was performed using the following conditions: 30 cycles of 98□°C for 2 min, 55□°C for 30□s and 68□°C for 0.5□min/kb. PCR products were examined for biallelic insertion using agarose gel electrophoresis. Initially, progenitor cells were produced by integrating OsTIR1 (CMV-OsTIR1) into the safe-harbor AAVS1 locus. We then co-transfected the donor template plasmid with Cas9 and sgRNAs that targeted the STOP codon of the LMNB1 or LMNB2 genes in HCT116 cells expressing OsTIR1. To generate cells with mAID in both genes, we simultaneously transfected the two donor templates with two sgRNA that target both genes. Cells expressing mClover for each protein target (LMNB1, LMNB2, or LMNB1 & B2) were sorted and grown as single cell colonies. We PCR-screened 179 colonies in total to identify clones with homozygous LAMINB1-mAID (5+/40 colonies), LAMIN B2-mAID (5+/55 colonies) as well as clones that had homozygous mAID in both genes (3+/84 colonies).

### Primers and sgRNA

Appropriate primers were designed to check the insertion by PCR. For creating stable cell lines, primer sets detected both the wild-type (WT) (1-1.5 kb) and inserted alleles (1-1.5 kb plus the size of the insertion), and another primer set to detect only the inserted allele. The first primer set was designed outside of the homology arms. Primers and sgRNA sequences used to create all AID cell lines are listed in **Supplementary Tables 1 – 3**.

### Plasmids for AID Cell lines

The donor construct contained the AID domain fused to mClover and an intervening T2A site with a hygromycin resistance marker. The construct was flanked by 50-base pair homology arms corresponding to the last exon region of the sgRNA recognition sequence and Cas9 cleavage site. To identify a CRISPR–Cas9 targeting site, we chose an appropriate sequence within 50 bp upstream or downstream from the stop codon. The following target finder sites were used to construct the CRISPR–Cas9 plasmid: IDT custom Alt-R guide design and WEG CRISPR finder. Construction of the CRISPR–Cas9 plasmid and donor plasmids have been previously described ^33^. The AAVS1 T2 CRISPR in pX330 plasmid (Addgene plasmid # 72833; http://n2t.net/addgene:72833; RRID: Addgene_72833) is based on pX330-U6-Chimeric_BB-CBh-hSpCas9 from Dr. Feng Zhang (Addgene #42230) (Cong et al, Science, 2013) and was a gift from Masato Kanemaki. The AAVS1 target sequence is described in Mali et al (Mali et al, Science, 2013). pMK232 (CMV-OsTIR1-PURO) was a gift from Masato Kanemaki (Addgene plasmid # 72834; http://n2t.net/addgene:72834; RRID: Addgene 72834). pMK364 (CMV-OsTIR1-loxP-PURO-loxP) was a gift from Masato Kanemaki (Addgene plasmid # 121184; http://n2t.net/addgene:121184; RRID: Addgene_121184). pMK290 (mAID-mClover-Hygro) was a gift from Masato Kanemaki (Addgene plasmid # 72828; http://n2t.net/addgene:72828; RRID: Addgene_72828).

### Auxin Treatment

For auxin treatment, HCT116^LMN(B1&B2)-AID^ cells were plated at 50,000 cells per well of a 6-well plate (Cellvis, P12-1.5H-N). To induce expression of OsTIR1, 2 μg/ml of doxycycline (Fisher Scientific, #10592-13-9) was added to cells 24 hours prior to auxin treatment. For live-cell flow cytometry, western blots, RT-qPCR, RNA-seq, and *in situ* Hi-C experiments, 1000 μM 3-Indoleacetic acid (IAA, Sigma Aldrich, #12886) was solubilized in 100% EtOH before each treatment as a fresh solution. For live-cell confocal microscopy, fixed-cell flow cytometry, fixed-cell immunofluorescence, CRISPR-Sirius fluorescent imaging, and Dual PWS experiments, 1000 μM Indole-3-acetic acid sodium salt (IAA, Sigma Aldrich, #6505-45-9) was solubilized in RNase-free water (Fisher Scientific, #10-977-015) before each treatment as a fresh solution and added to HCT116^LMN(B1&B2)-AID^ cells. Optimal auxin treatment time was determined based on the results from western blot, immunofluorescence, and flow cytometry experiments **(Supplementary Fig. 5)**.

### Fixed-Cell Flow Cytometry (FACS) Analysis

Flow cytometry analysis for HCT116^LMNB1-AID^, HCT116^LMNB2-AID^ and HCT116^LMN(B1&B2)-AID^ cells for AID system verification and nuclear morphology experiments was performed on the Amnis ImageStreamXTM, located at the University of Virginia Flow Cytometry Core Facility in Charlottesville, VA. To assess the degree of apoptosis induced by auxin treatment, we used the Annexin V APC Kit (Cayman Chemical, #601410) and followed the manufacturer’s protocol. Flow cytometry analysis for HCT116^LMN(B1&B2)-AID^ cells to determine proper auxin treatment concentration was performed on a BD LSRFortessa Cell Analyzer FACSymphony S6 SORP system, located at the Robert H. Lurie Comprehensive Cancer Center Flow Cytometry Core Facility at Northwestern University in Evanston, IL. For all FACS analysis the same protocol was used. After 24 hours of doxycycline treatment followed by auxin treatment, cells were harvested and fixed. Briefly, cells were washed with DPBS (Gibco, #14190-144), trypsinized (Gibco, #25200-056), neutralized with media, and then centrifuged at 500 x g for 5 minutes. Cells were then resuspended in 500 μL of 4% PFA and DPBS and fixed for 10 minutes at room temperature, followed by centrifugation and resuspension in cold FACS buffer (DPBS with 1% of BSA and 2mM EDTA added) at 4°C until analysis could be performed the following day. Data were analyzed using FlowJo software.

### Protein Detection & Antibodies

HCT116^LMNB1-AID^, HCT116^LMNB2-AID^ and HCT116^LMN(B1&B2)-AID^ cells were lysed using Radio Immuno Precipitation Assay (RIPA) buffer (Sigma-Aldrich, #R0278) with protease inhibitor added (Sigma-Aldrich, #P8340). Cell lysates were quantified with a standard Bradford assay using the Protein Assay Dye Concentrate (BioRad, #500-0006) and BSA as a control. Heat denatured protein samples were resolved on a 4-12% bis-tris gradient gel, transferred to a PVDF membrane using the Life Technologies Invitrogen iBlot Dry Transfer System (Thermo Fisher Scientific, IB1001) (20V for 7 minutes), and blocked in 5% nonfat dried milk (BioRad, #120-6404) in 1x TBST. Whole-cell lysates were blotted against the following primary antibodies: Lamin B1 (Cell Signaling, #13435, dilution 1:1000), Lamin B2 (Cell Signaling, #12255, dilution 1:1000), and alpha-tubulin (Thermo Fischer Scientific, #62204, dilution 1:2000). The following secondary antibodies were used: anti-rabbit IgG HRP (Promega, #W4018). Blots were incubated with the primary antibody overnight at 4□°C, followed by incubation with the secondary antibodies for 1 hour at room temperature. To develop blots for protein detection, chemiluminescent substrates were used (Thermo Fischer Scientific, #32106).

### Quantitative Real-Time PCR & RNA Isolation

Total RNA from transfected cells was harvested using the RNeasy Plus Mini Kit (Qiagen, #74134) following the manufacturer’s protocol. 1 mg of RNA was converted to cDNA using the Applied Biosystems High-Capacity RNA-to-cDNA Kit (Thermo Fisher Scientific, #4387406). The GAPDH or HPRT-1 gene was used as an internal control for analysis. RT-qPCR was performed on a StepOnePlus Applied Biosystems instrument with SYBR Green. RNA quantity was measured using the Nanodrop 2000 Spectrophotometer at 260nm.

### Fixed-Cell Immunofluorescence

HCT116^LMN(B1)-AID^ cells, HCT116^LMN(B2)-AID^ cells, or HCT116^LMN(B1&B2)-AID^ cells at a low passage (<P10) were plated at 100,000 cells per well of a 6-well glass-bottom plate (Cellvis, #P06-1.5H-N). Following auxin treatment, cells were washed twice with 1x Phosphate Buffered Saline (PBS) (Gibco, #10010031). Cells were fixed with 4% paraformaldehyde (PFA) (Electron Microscopy Sciences, #15710) for 10 minutes at room temperature, followed by washing with PBS 3 times for 5 minutes each. Cells were permeabilized using 0.2% TritonX-100 (10%) (Sigma-Aldrich, #93443) in 1x PBS, followed by another wash with 1x PBS for 3 times for 5 minutes each. Cells were blocked using 3% BSA (Sigma-Aldrich, #A7906) in PBST (Tween-20 in 1x PBS) (Sigma-Aldrich, #P9416) at room temperature. The following primary antibodies were added overnight at 4□°C: lamin B1 (Abcam, #ab16048, dilution 1:1000), lamin B2 (Abcam, #ab155319, dilution 1:1000. Cells were washed with 1x PBS 3 times for 5 minutes each. The following secondary antibody was added for 1 hour at room temperature: Invitrogen Goat anti-Rabbit IgG (H+L) Highly Cross-Adsorbed Secondary Antibody, Alexa Fluor 646 (Thermo Fisher Scientific, #A-21245). Cells were washed with 1x PBS 3 times for 5 minutes each. Finally, cells were stained with DAPI (Thermo Fisher Scientific, #62248, diluted to 0.5 μg/mL in 1x PBS) for 10 minutes at room temperature. Prior to imaging, cells were washed with 1x PBS twice for 5 minutes each.

### Preparation of Hi-C Libraries for *in situ* Hi-C

*In-situ* Hi-C was performed as previously described ^2^. Briefly, 2-5 million cells at 80% confluence were detached and pelleted by centrifugation at 300xG for 5 minutes. Cells were resuspended in fresh medium at a concentration of 1E6 cells per 1 mL media. In a fume hood, 1% formaldehyde was used to crosslink cells, with 10 minutes of incubation at room temperature with mixing. 2.5M glycine solution was added to a final concentration of 0.2M to quench the reaction, and the reaction was incubated at room temperature for 5 minutes with gentle rocking. Samples were centrifuged at 300xG for 5 minutes at 4°C. Cells were resuspended in 1mL of ice-cold PBS and spun at 300xG for 5 minutes at 4°C. Cell pellets were flash-frozen in liquid nitrogen and either stored at −80°C or used immediately for lysis and restriction digest. Nuclei were permeabilized. 250 μL of Hi-C lysis buffer (10mM Tris-HCl pH 8.0, 10mM NaCl, 0.2% Igepal CA930 (Sigma-Aldrich, #I3021)) with 50 μL of protease inhibitors (Sigma Aldrich, #P8340) was added to each crosslinked pellet of cells. After incubation on ice for >15 minutes, samples were centrifuged at 2500xG for 5 minutes and washed with 500 μL of ice-cold Hi-C lysis buffer. Pelleted nuclei were resuspended in 50 μL of 0.5% sodium dodecyl sulfate (SDS) (Sigma-Aldrich, #436143) and incubated at 62°C for 10 minutes. Next, 145 μL of water (Fischer Scientific, #10-977-015) and 25 μL of 10% Triton X-100 (Sigma Aldrich, #93443) were added to quench the SDS. Samples were mixed and incubated at 37°C for 15 minutes. DNA was digested with 100U of MboI restriction enzyme (New England Biolabs, #R0147) and 25 μL of 10X NEBuffer 2 (New England Biolabs, #B7002S). Chromatin was digested overnight at 37°C with rotation. Samples were incubated at 62°C for 20 minutes to inactivate MboI, and then cooled to room temperature. The ends of restriction fragments were labeled using biotinylated nucleotides (Thermo Fisher Scientific, #19524016) and ligated in a small volume (∼ 900 μL) using 10X NEB T4 DNA ligase buffer (New England Biolabs, #B0202) and DNA Polymerase I, Large (Klenow) Fragment (New England Biolabs, #M0202) after 1 hour of incubation at 37°C. Samples were mixed and incubated at room temperature for 4 hours prior to reversal of crosslinks. We added 50 μL of 20 mg/mL proteinase K (New England Biolabs, #P8102) and 120 μL of 10% SDS and incubated samples at 55°C for 30 minutes. Next, 130 μL of 5M sodium chloride was added and samples were incubated at 68°C overnight. Ligated DNA was purified and sheared to a length of ∼400 bp, as previously described ^2^ using a LE220-plus Focused-ultrasonicator (Covaris, #500569) and AMPure XP beads (Beckman Coulter, #A63881). DNA was quantified using the Qubit dsDNA High Sensitivity Assay Kit (Thermo Fisher Scientific, #Q33230) and undiluted DNA was run on a 2% agarose gel to verify successful size selection. Point ligation junctions were pulled down with 10 m/mL Dynabeads MyOne Steptavidin T1 beads (Thermo Fisher Scientific, #65601) and prepared for Illumina sequencing using Illumina primers and protocol (Illumina, 2007) as previously described ^2^. Paired-end sequencing was performed using the Illumina HiSeq 2000 OR 2500 platform. A no-ligation control was also used.

### Hi-C Data Processing and Analysis

Juicebox was used to visualize Hi-C contact maps^66^. All Hi-C data reported were produced using Illumina paired-end sequencing. We followed the Hi-C data processing pipeline that has previously been described ^2^. This pipeline uses the Burrows-Wheeler single end aligner (BWA)^67^ to map each read end separately to the hg19 reference genome, removes reads that map to the same fragment, removes duplicate or near-duplicate reads, and filters the remaining reads based on the mapping quality score. All analysis (i.e., aggregate peak analysis) and annotations (i.e., annotation of domains, assigning loci to subcompartments, and peaks) were performed as previously described ^2,68^. All contact matrices were KR-normalized with Juicer. Domains were annotated using TopDom.

### Dual PWS Imaging

Briefly, PWS measures the spectral interference signal resulting from internal light scattering originating from nuclear chromatin. This is related to variations in the refractive index distribution (Σ) (extracted by calculating the standard deviation of the spectral interference at each pixel), characterized by the chromatin packing scaling (*D*). *D* was calculated using maps of Σ, as previously described ^40,50,52,53^. Measurements were normalized by the reflectance of the glass medium interface (i.e., to an independent reference measurement acquired in a region lacking cells on the dish). This allows us to obtain the interference signal directly related to refractive index (RI) fluctuations within the cell. Although it is a diffraction-limited imaging modality, PWS can measure chromatin density variations because the RI is proportional to the local density of macromolecules (e.g., DNA, RNA, proteins). Therefore, the standard deviation of the RI (Σ) is proportional to nanoscale density variations and can be used to characterize packing scaling behavior of chromatin domains with length scale sensitivity around 20 – 200 nm, depending on sample thickness and height. Changes in *D* resulting from each condition are quantified by averaging over nearly 2000 cells, taken across 3 technical replicates. Live-cell PWS measurements obtained using a commercial inverted microscope (Leica, DMIRB) using a Hamamatsu Image-EM charge-coupled device (CCD) camera (C9100-13) coupled to a liquid crystal tunable filter (LCTF, CRi VariSpec) to acquire monochromatic, spectrally resolved images ranging from 500-700 nm at 2-nm intervals as previously described ^47,50,51^. Broadband illumination is provided by a broad-spectrum white light LED source (Xcite-120 LED, Excelitas). The system is equipped with a long pass filter (Semrock BLP01-405R-25) and a 63x oil immersion objective (Leica HCX PL APO). Cells were imaged under physiological conditions (37°C and 5% CO_2_) using a stage top incubator (In vivo Scientific; Stage Top Systems). All cells were given at least 24 hours to re-adhere before treatment (for treated cells) and imaging.

### Dynamic PWS Measurements

Dynamic PWS measurements were obtained as previously described.^50^ Briefly, dynamics measurements (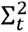, fractional moving mass (*m*_f)_, and diffusion) are collected by acquiring multiple backscattered wide-field images at a single wavelength (550□nm) over time (acquisition time), to produce a three-dimensional image cube, where 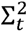 is temporal interference and t is time. Diffusion is extracted by calculating the decay rate of the autocorrelation of the temporal interference as previously described.^50^ The fractional moving mass is calculated by normalizing the variance of 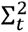 at each pixel. Using the equations and parameters supplied and explained in detail in the supplementary information of our recent publication ^50^, the fractional moving mass is obtained by using the following equation to normalize 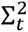 by ρ_0_, the density of a typical macromolecular cluster:

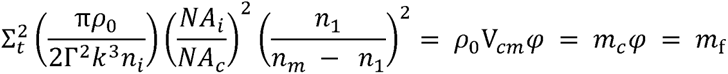

With this normalization, 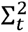 is equivalent to *m*_f_, which measures the mass moving within the sample. This value is calculated from the product of the mass of the typical moving cluster (*m_C_*) and the volume fraction of mobile mass (φ). *m_C_* is obtained by *m_C_* = V*_Cm_ ρ*_0_, where V*_Cm_* is the volume of the typical moving macromolecular cluster. To calculate this normalization, we approximate *n_m_* = 1.43 as the refractive index (RI) of a nucleosome, *n*_1_ = 1.37 as the RI of a nucleus, *n_i_* = 1.518 as the refractive index of the immersion oil, and *ρ* _0_ = 0.55 g *cm*^-3^ as the dry density of a nucleosome. Additionally, *k* = 1.57E5 cm^-1^ is the scalar wavenumber of the illumination light, and Γ is a Fresnel intensity coefficient for normal incidence. *NA_C_* = 1.49 is the numerical aperture (NA) of collection and *NA_i_* = 0.52 is the NA of illumination. As stated previously^50^, 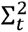 is sensitive to instrument parameters such as the depth of field, substrate refractive index, etc. These dependencies are removed through normalization with the proper pre-factor calculated above for obtaining biological measurements. It should also be noted that backscattered intensity is prone to errors along the transverse direction^50^. Due to these variations, these parameters are more accurate when calculating the expected value over each pixel.

### Regional PWS Analysis

We used PWS to calculate *D* values via measuring the variations in spectral light interference resulting from light scattering due to heterogeneities in chromatin density as previously described ^52^. The same cells used to analyze average chromatin packing scaling in whole nuclear regions were used for regional chromatin packing scaling analysis. For the periphery characterization, individual nuclei were segmented into 6 ribbons of 260 nm width each using MATLAB. The remaining region at the center of the nucleus was classified as the center. We calculated the average *D* for each pixel, followed by averaging all these values to estimate the average *D* in each region.

### Confocal Imaging

For CRISPR-Sirius, when MCP-Halo was added to cells for fluorescent imaging, HaloTag-JF646 was added to the cells at 10 μM 24□h before imaging and incubated overnight at 37°C and 5% CO_2_. On the day of imaging, live cells were washed three times with DPBS (Gibco, #14190-144) and further incubated with phenol-red free media (Cytiva, #SH30270.01). The optical instrument was built on a commercial inverted microscope (Eclipse Ti-U with the perfect focus system, Nikon). Images of live cells were collected using a 100× objective and sent to an electron-multiplying CCD (iXon Ultra 888, Andor). A 637 nm laser (Obis, Coherent) was co-illuminated through a 100x/ 1.49 NA (numerical aperture) oil objective lens (SR APO TIRF, Nikon) with an average power at the sample of 3 to 10 kW/cm^3^. The microscope stage incubation chamber was maintained at 37 °C and supplemented with 5% CO_2_. For single image acquisition, at least 50 frames were taken at 30 ms exposure time and a gain of 150. Z-stack images were acquired at 0.024 µm per step for a total of 401-701 frames depending on the size of the nuclei being imaged. ^47,50,51^ Images of fixed-cells previously transfected with CRISPR-Sirius plasmids were imaged using the Nikon SoRa Spinning Disk confocal microscope equipped with a Hamamatsu ORCA-Fusion Digital CMOS camera. Images were collected using a 60x/ 1.42 NA oil-immersion objective mounted with a 2.8x magnifier. mClover was excited with a 488 nm laser, HaloTag-JF646 was excited with a 640 nm laser, and DAPI was excited with a 405 nm laser. Imaging data were acquired by Nikon acquisition software.

### Lentivirus Packaging

HEK293T cells were used to produce lentiviral particles using FuGENE HD Transfection Reagent (Promega, #E2311) following the manufacturer’s protocol. Briefly, one day before transfection, HEK293T cells at low passage (<P10) were plated at 100,000 cells per well of a 12-well plate (Cellvis, P12-1.5H-N). At the time of transfection, cells reached a confluency of 70-80%. For lentivirus packaging, a master mix of DNA was prepared in reduced serum media (OptiMEM, Gibco, #31985-070). This master mix contained the lentiviral packaging plasmid pCMV-VSV-G (a gift from Bob Weinberg, Addgene plasmid # 8454) and pCMV-dR8.2 (a gift from Bob Weinberg, Addgene plasmid # 8455). For packaging each virus, the following amounts of each plasmid were mixed: 0.5 μg transfer vector + 0.45 μg pCMV-dR8.2 + 0.05 μg pCMV-VSV-G. Media was changed 24 hours post-transfection to fresh DMEM. Lentiviral particles were harvested 60 hours after transfection. The viral supernatants were filtered using a 33 mm diameter sterile syringe filter with a 0.45 µm pore size hydrophilic PVDF membrane (Millipore Sigma, SLHVR33RS) and added to HEK293T cells. The virus was immediately used or stored at −80□°C. Polybrene (8 μg/mL; Sigma-Aldrich) was supplemented to enhance transduction efficiency.

### CRISPR-Sirius Labelling

Plasmids were obtained from Addgene as bacterial stabs and streaked onto LB-ampicillin plates. Upon overnight growth and single colony selection, a single colony was inoculated into LB-ampicillin liquid culture overnight. Plasmid DNA isolation was performed using QIAprep Spin Miniprep kit (Qiagen, # 27104) following the manufacturer’s protocol. pHAGE-TO-dCas9-P2A-HSA (Addgene plasmid # 121936; http://n2t.net/addgene:121936; RRID: Addgene_121936), pHAGE-EFS-MCP-HALOnls (Addgene plasmid # 121937; http://n2t.net/addgene:121937; RRID: Addgene_121937), and pPUR-hU6-sgRNA-Sirius-8XMS2 (Addgene plasmid # 121942; http://n2t.net/addgene:121942; RRID: Addgene_121942) were gifts from Thoru Pederson.

### CRISPR-Sirius Transduction

For live-cell CRISPR-Sirius, HCT116^LMN(B1&B2)-AID^ cells were transfected with CRISPR-dCas9 and donor plasmids using FuGENE HD Transfection Reagent (Promega, #E2311) following the manufacturer’s protocol. Briefly, cells at low passage (<P10) were plated at 100,000 cells per well of a 6-well glass-bottom plate (Cellvis, P06-1.5H-N). 24 hours after plating, 50 μL dCas9, 50 μL MCP-HALOnls, and 100 μL sgRNA lentiviral particles were added to each well. 24 hours after transduction, lentiviral particles were removed by replacing media.

### CRISPR-Sirius Transfection

For additional quantification of foci and distances of foci to the nuclear periphery, HCT116^LMN(B1&B2)-AID^ cells were co-transfected with 200 ng MCP-HaloTag, 400 ng of dCas9 plasmid DNA, and 2 µg of plasmid DNA for the desired guide RNAs using Lipofectamine LTX and Plus Reagent. Cells were incubated for 24 hours prior to overnight staining with HaloTag-JF646 before fixation and imaging.

### Data and Image Analysis

We used GraphPad Prism 9.3.1 or Excel for statistical analysis and for making all boxplots. Flow cytometric analysis (FACS) data were analyzed using FlowJo software version 10.6.1. To localize the fluorescent puncta, we used ImageJ software to first generate max projections of the Z-stack images/50 frame single layer acquisition. Max projected images were then background subtracted using the standard rolling ball algorithm with a radius of 12.0. Processed images were then input into the Thunder-STORM ImageJ plugin with a peak intensity threshold coefficient of 5.0 to locate the fluorescent puncta associated with the CRISPR Sirius tagged loci. Puncta coordinates were saved in Microsoft Excel as a .csv file to be input to the Python algorithm. The processed image was then fed into a Python computer vision algorithm that segmented out the nucleus and determined the coordinates for the nuclear periphery as well as the centroid of the nucleus. To quantify the spatial distance to the nuclear periphery, only pairs of loci lying in the same foci plane were analyzed. Distances were measured from each fluorescent puncta to the nearest nuclear periphery coordinate as well as the centroid of the segmented nucleus. To account for differences in cell area, the distance from each puncta to the centroid was divided by the radius, assuming a circular area. This was further verified by dividing the distance by the nuclear area. To detect loci numbers, maximum intensity projection of Z-series images was performed.

### Coefficient of Variation Analysis

To assess chromatin compaction through the Coefficient of Variation (CV) analysis, DAPI-stained cells (see section Fixed-cell immunofluorescence) treated with Auxin (see section Auxin treatment) were imaged on a Nikon SoRa Spinning Disk confocal microscope (see section Confocal imaging). Following a published workflow^54^, we used ImageJ to create masks of each nucleus. The coefficient of variation of individual nuclei was calculated in MATLAB, with CV = σ/μ, where σ represents the standard deviation of the intensity values and μ representing the mean value of intensity of the nucleus.

### RNA-Seq Library Preparation

Total RNA extraction was performed on samples from colon carcinoma epithelial HCT116 cells and HCT116^LMN(B1&B2)-AID^ cells using the RNeasy Plus Mini Kit (Qiagen, #74134) following the manufacturer’s protocol. The conditions for these samples were control, 12-hour auxin, 48-hour auxin, and 48-hour auxin with 6 days of removal by changing cell culture media. These samples were collected with three biological replicates per condition. RNA-Seq libraries were prepared using the NEBNext Ultra Directional RNA Library Prep Kit for Illumina (New England BioLabs, #E7760), according to the company’s instruction. Library quality was measured using the Qubit 2.0 and Bioanalyzer.

### RNA-Seq Data Analysis

Bulk mRNA sequencing was conducted in The Genome Analysis and Technology Core in Charlottesville, VA at the University of Virginia. Paired-end reads were acquired using NextSeq 2000 (75bp) system on high throughput mode. Reads were aligned to the hg19 genome using HISAT2 and quantified using StringTie. Read counts were normalized and compared for differential gene expression using the DESeq2 package in R. Heatmaps were generated using the pheatmap package. Other plots were generated using the ggplot2 package. We used Metascape (https://metascape.org/gp/index.html#/main/step1) to perform pathway enrichment analysis. The publicly available hg19 DamID track was downloaded from the UCSC genome browser database. We used bedtools to compare gene coordinates with the DamID LAD coordinates. We specifically used the “reldist”, “closest”, and “coverage” options of bedtools.

## QUANTIFICATION AND STATISTICAL ANALYSIS

Experimental data are presented as the mean ± SD of three independent experiments unless otherwise stated. Statistical analysis was performed using two-sided Student’s t-test for comparing two sets of data with normal distribution assumed, unless stated otherwise. For datasets with a Gaussian distribution, parametric tests were applied. For datasets with no Gaussian distribution, non-parametric tests were applied. In all cases two-tailed test were run and multiple comparison corrections were applied for datasets with more than two groups and multiple comparisons. Statistical analysis for PWS, coefficient of variation, and CRISPR-Sirius plots has been performed in GraphPad Prism. A *P* value of < 0.05 was considered significant. Statistical significance levels are denoted as follows: ns; \**P*<0.05; \*\**P*<0.01; \*\*\**P*<0.001; \*\*\*\**P*<0.0001. Sample numbers (# of nuclei, n) and the number of replicates (N) is indicated in figure legends.

## DECLARATIONS

## ETHICS APPROVAL AND CONSENT TO PARTICIPATE

Not applicable.

## CONSENT FOR PUBLICATION

Not applicable.

## AVAILABILITY OF DATA AND MATERIALS

The original source code during the current study is available in the Backman Lab/ Lamin-Project GitHub repository (https://github.com/BackmanLab/Lamin-Project). Datasets used and/or analyzed during the current study are available from the corresponding author on reasonable request. Raw data is available from the corresponding authors upon reasonable request. The cell lines have been authenticated and are available upon request. Further information and requests for resources and reagents should be directed to and will be fulfilled by the lead contact, Vadim Backman.

## AUTHORS’ CONTRIBUTIONS

E.P., X.W, J.Y., and N.A., and F.S. conducted the experiments; E.P. wrote the paper; L.C and J.Y. conducted RNA-seq analysis; E.P and N.A. assisted with PWS imaging and performed analysis; S.R., L.C., and L.A. conducted *in* situ Hi-C analysis; A.D. wrote the code for analyzing regional chromatin packing scaling and dynamics; V.A., L.C., and L.A. edited the manuscript.

## Supporting information

Key Resources Table

## ACKNOWLEDGEMENTS

We thank S. Gladstein, N. Anthony, S. Jain, J. Frederick, R. Virk, and A. Shim for their helpful discussions during the experimental design and manuscript preparation. We also thank B. Keane for proofreading the manuscript.

## INCLUSION AND DIVERSITY

One or more of the authors of this paper self-identifies as an underrepresented minority.

---

**Supplementary Figure 1.**
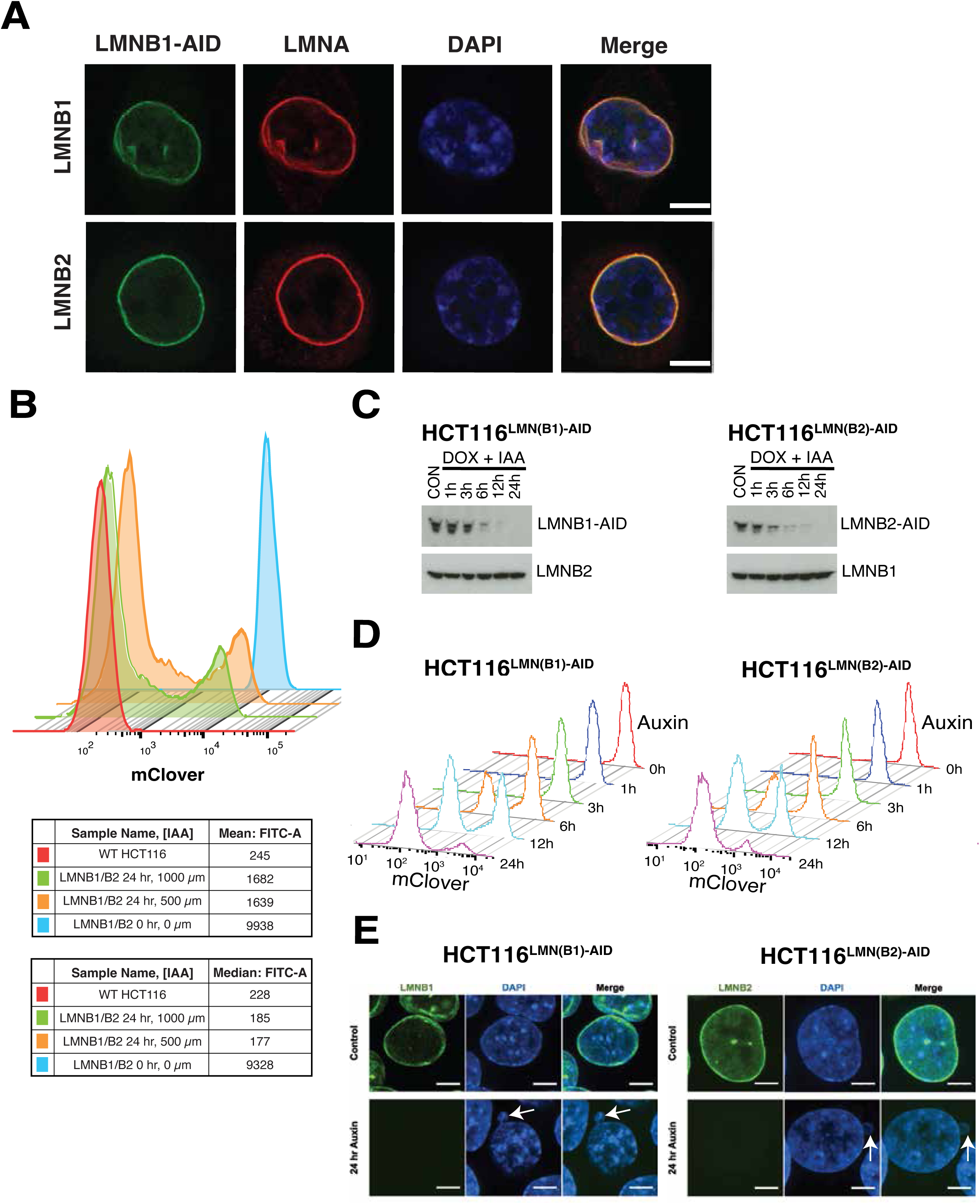

**Supplementary Figure 2.**
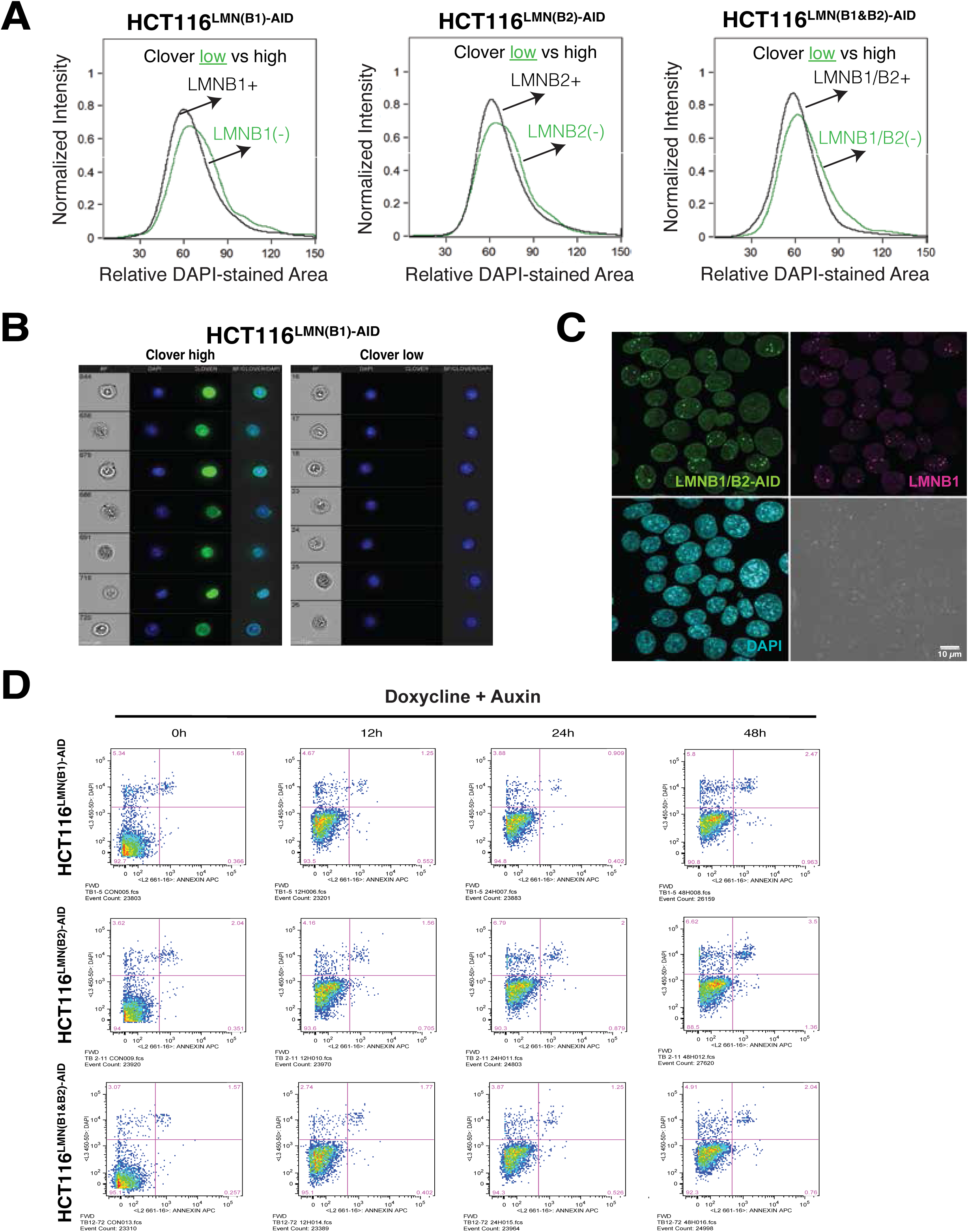

**Supplementary Figure 3.**
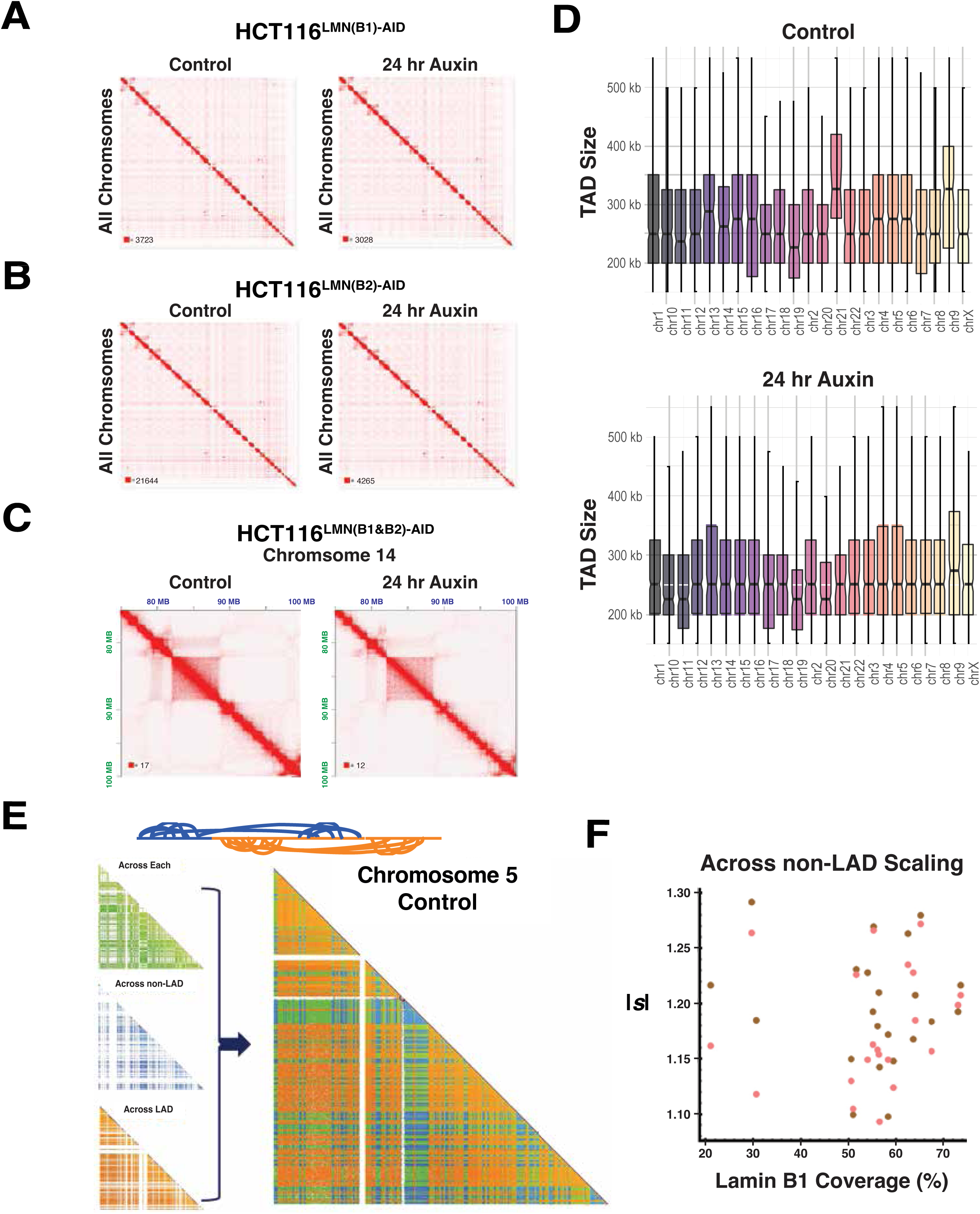

**Supplementary Figure 4.**
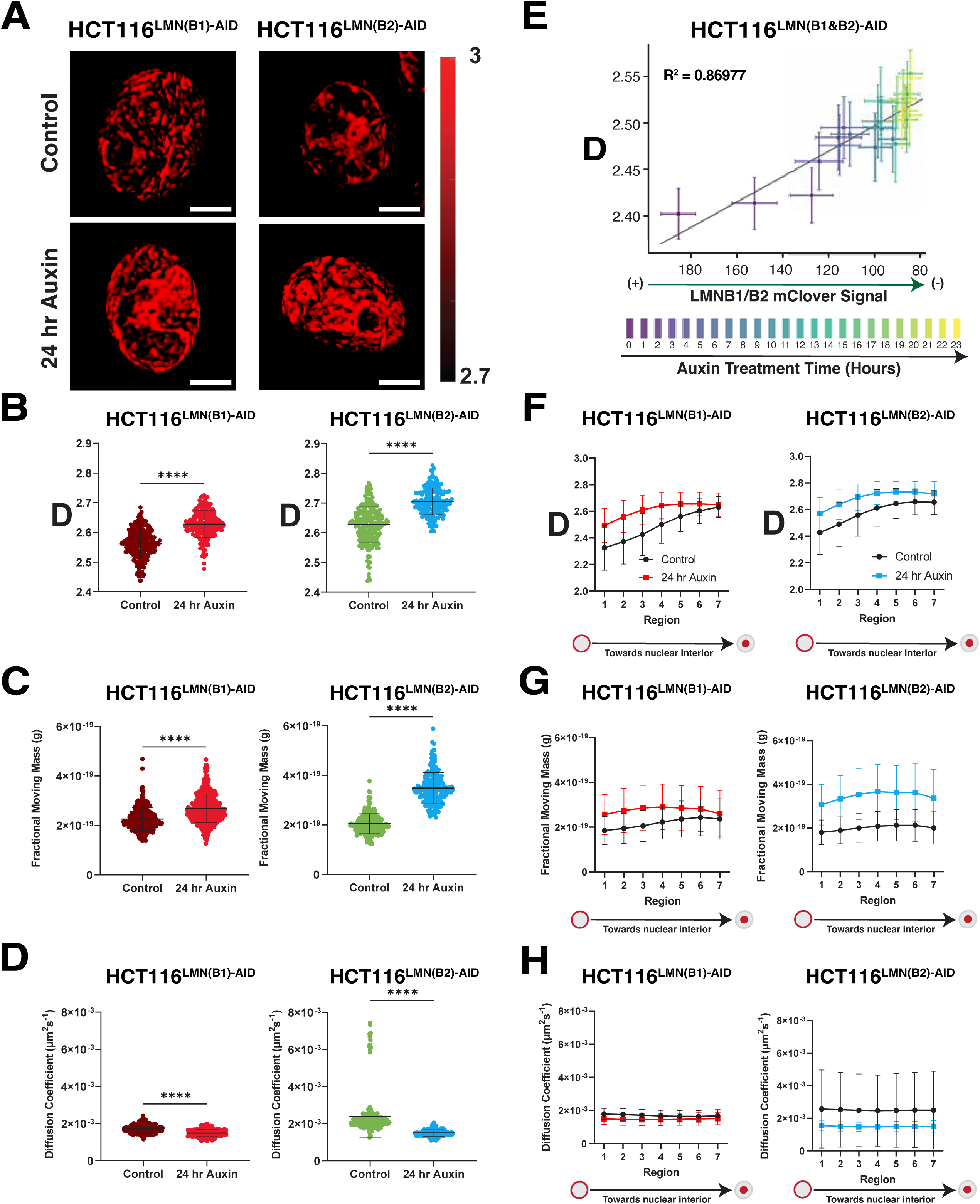

**Supplementary Figure 5.**
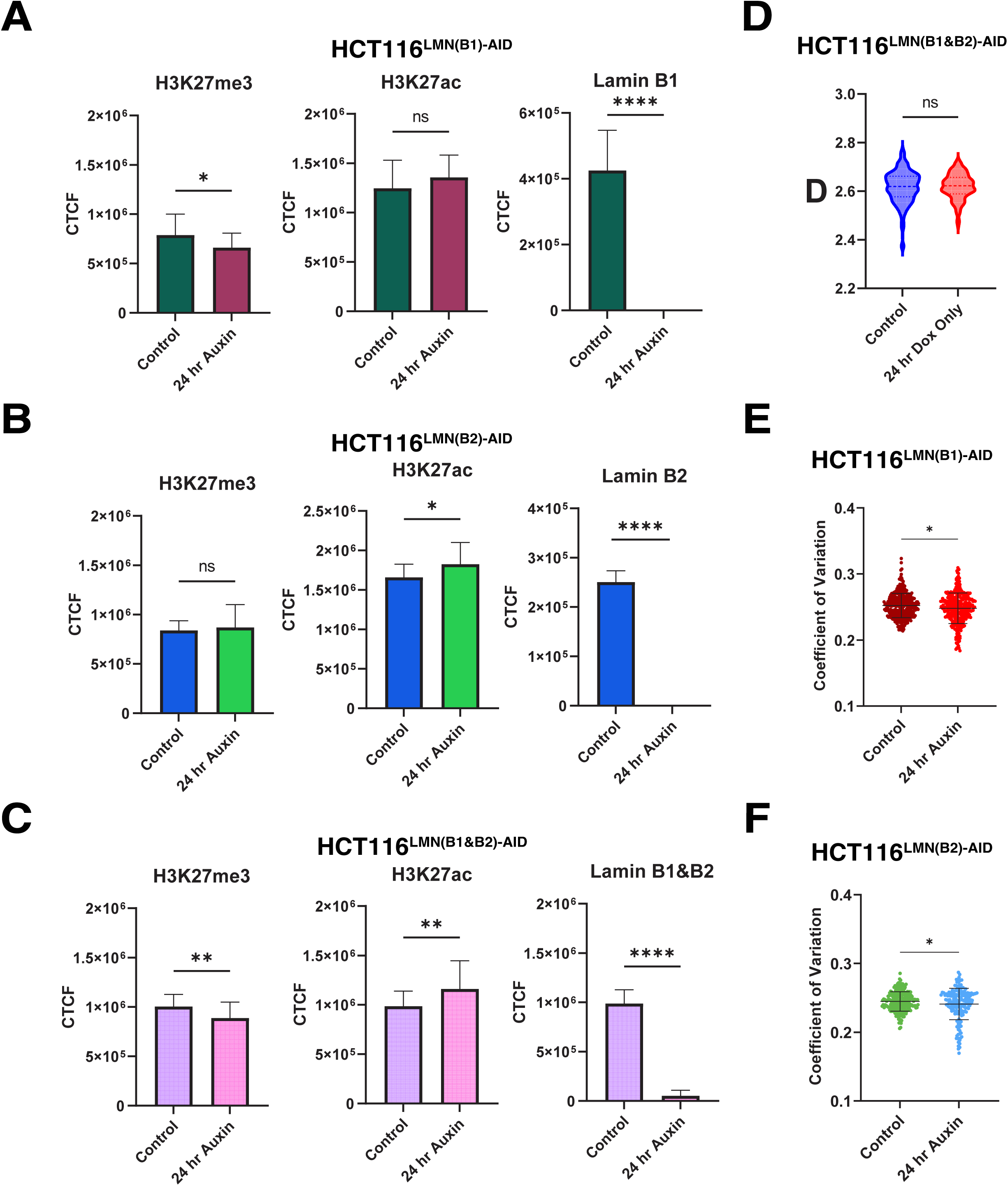

**Supplementary Figure 6.**
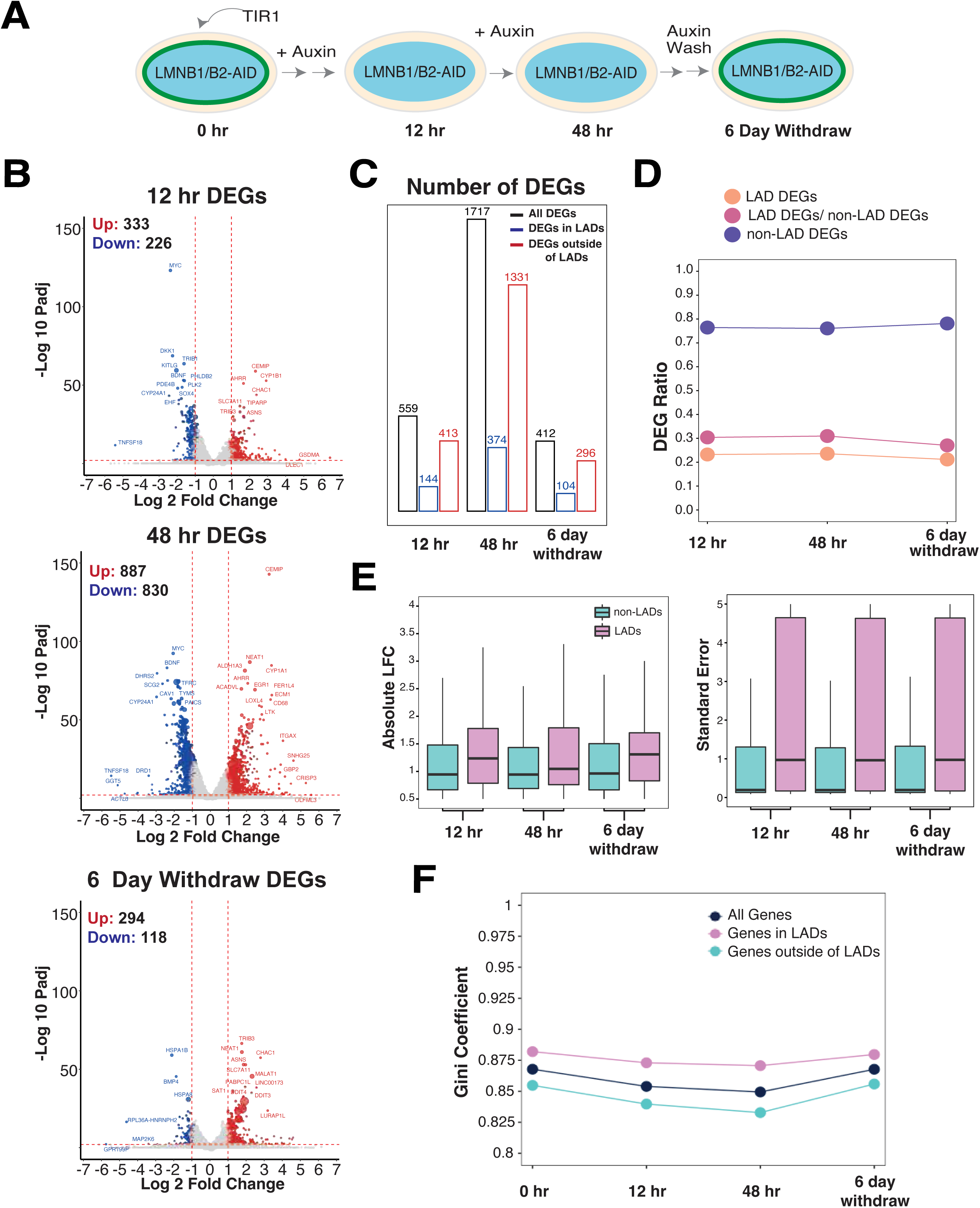

