## Supplementary Information for "Depletion of lamins B1 and B2 alters chromatin mobility and induces differential gene expression by a mesoscale-motion dependent mechanism"

**SUPPLEMENTARY FIGURES**

**Supplementary Figure 1:**

**Addition of the mAID tag does not change localization of Lamin A, and auxin treatment results in reduced Lamin B proteins and increased nuclear blebbing.**

(A) The nuclear localization of LMNB1-AID (green) and LMNA (red) in HCT116LMN(B1)-AID cells was examined by immunofluorescence. Cells were stained with DAPI to indicate the nuclear region. Scale bar = 10 μm. N = 3, data was obtained from three independent biological replicates.
(B) According to the flow cytometry results, 1000 µM of 24-hour auxin treatment is the optimal concentration for maximal degradation of LMNB1 and LMNB2 in fixed HCT116^LMN(B1&B2)-AID^ cells. At least 20,000 events were recorded during the experiment.
(C) Western blots reveal that AID-tagged LMNB1 and LMNB2 are degraded within 24 hours of auxin treatment. Within this time frame, the target protein expression is no longer detectable in HCT116LMN(B1)-AID cells and HCT116LMN(B2)-AID cells.
(D) Single cell-level degradation kinetics demonstrate a loss of B-type lamin proteins, indicated by the loss of the mClover signal. Within 24 hours of auxin treatment, the majority of mClover signal is lost in HCT116^LMN(B1)-AID^ cells and HCT116^LMN(B2)-AID^ cells.
(E) Immunofluorescence shows the presence of nuclear blebbing in B-type lamin-deficient cells.

**Supplementary Figure 2:**

**Degradation of B-type lamins increases nuclear area but does not induce apoptosis.**

(A) Flow cytometric analysis reveals that in HCT116^LMNB1-AID^, HCT116^LMNB2-AID^ and HCT116^LMN(B1&B2)-AID^ cells, 24 hours of auxin treatment (Clover negative/ low) shifts the intensity of mClover to be nearly undetectable. Auxin treatment increases the nuclear area (DAPI +).
(B) Representative images from ImageStreamX show HCT116^LMNB1-AID^ cells after 8 hours of auxin treatment with depleted mClover signal and increased nuclear area.
(C) Confocal fluorescence microscopy in fixed HCT116 LMN^(B1&B2)-AID^ cells reveal lamin-specific puncta that are spherical and discrete in organization. AID-tagged LMNB1/B2 is shown in green while endogenous LMNB1 is shown in magenta. Aggregates of high lamin expression are shown. Cells were stained with DAPI to indicate the nuclear region. The brightfield image is shown on the bottom right. Scale bar ~ 10 μm. N = 3; data was obtained from three independent biological replicates.
(D) Auxin treatment to induce acute depletion of B-type lamins in HCT116^LMN(B1)-AID^ cells, HCT116^LMN(B2)-AID^ cells, and HCT116^LMN(B1&B2)-AID^ cells does not substantially induce cell death. Viable, in-tact cells are present in the bottom left quadrant, dead cells are shown in the top right quadrant, and apoptotic cells are shown in the bottom right quadrant in the plots.

**Supplementary Figure 3:**

**The mesoscale structure of chromatin is overall preserved upon B-type lamin degradation.**

(A) The normalized Hi-C trans-interaction matrices for all chromosomes in the control and 24-hour auxin treatment are shown for HCT116^LMN(B1)-AID^ cells.
(B) The normalized Hi-C trans-interaction matrices for all chromosomes in the control and 24-hour auxin treatment are shown for HCT116^LMN(B2)-AID^ cells.
(C) The normalized Hi-C interaction matrices for HCT116^LMN(B12&B2)-AID^ cells for chromosome 14 and 50 kb resolution is shown from 75 Mb to 100 Mb.
(D) TAD sizes for each chromosome are shown for HCT116^LMN(B1&B2)-AID^ cells in the control and 24-hour auxin treatment conditions.
(E) The schematic demonstrates contacts across LAD segments and non-LAD segments for a representative chromosome. The Hi-C trans-interaction matrix demonstrates contacts across LAD segments, non-LAD segments and contacts across both segments for chromosome 5 (control condition).
(F) The scatter plot for the relationship between lamin B1 coverage and |*s|* in non-LAD segments is shown.

**Supplementary Figure 4:**

**Dual Partial Wave Spectroscopic (PWS) microscopy reveals differential higher-order chromatin structure and dynamics upon loss of either lamin B1 or lamin B2.**

(A) Representative images of HCT116^LMN(B1)-AID^ cells and HCT116^LMN(B2)-AID^ cells obtained from PWS microscopy, a label-free, high throughput interference-based imaging technique are shown for both the control and 24-hour auxin treatment conditions.
(B) Violin plots demonstrate that chromatin packing scaling (D) is significantly increased in HCT116^LMN(B1)-AID^ cells and HCT116^LMN(B2)-AID^ cells.
(C) Violin plots demonstrate that fractional moving mass is significantly increased in HCT116^LMN(B1)-AID^ cells and HCT116^LMN(B2)-AID^ cells upon 24 hours of auxin treatment.
(D) Violin plots demonstrate that the diffusion coefficient is significantly decreased in HCT116^LMN(B1)-AID^ cells and HCT116^LMN(B2)-AID^ cells upon 24 hours of auxin treatment. (B-D). The truncated violin plots extend from the minimum to the maximum value. The line in the middle of each plot is the median value of the distribution, and the lines above and below are the third and first quartiles, respectively. Data was obtained from three technical replicates for each condition. ****P<0.0001. Welch’s correction was applied.
(E) A PWS 24-hour auxin treatment time course shows the relationship between *D* and mClover signal over 24 hours of auxin treatment for HCT116^LMN(B1&B2)-AID^ cells. Data was obtained from three technical replicates for each condition (N=3, *n = 634*). ****P<0.0001.
(F) Regional PWS measurements of *D* in HCT116^LMN(B1)-AID^ cells and HCT116^LMN(B2)-AID^ cells are shown, along with the change in *D* within each region.
(G) Regional PWS measurements of fractional moving mass for HCT116^LMN(B1)-AID^ cells and HCT116^LMN(B2)-AID^ cells are shown, along with the change in fractional moving mass within each region.
(H) Regional PWS measurements of the diffusion coefficient for HCT116^LMN(B1)-AID^ cells and HCT116^LMN(B2)-AID^ cells are shown, along with the change in the diffusion coefficient within each region.

**Supplementary Figure 5:**

**Loss of B-type lamins results in the shifted distribution of repressive marks towards the nuclear interior and chromatin decompaction.**

(A) The bar plots show H3K27me3, H3K27ac, and Lamin B1 corrected total cell fluorescence in HCT116^LMN(B1)-AID^ cells for the control (untreated) and 24-hour auxin treatment conditions. Data was obtained from three technical replicates for each condition (N=3, Control (*n* = 281), 24-hour auxin (*n* = 193)). ****P<0.0001.
(B) The bar plots show H3K27me3, H3K27ac, and Lamin B1 corrected total cell fluorescence in HCT116^LMN(B2)-AID^ cells for the control (untreated) and 24-hour auxin treatment conditions. Data was obtained from three technical replicates for each condition (N=3, Control (*n* = 249), 24-hour auxin (*n* = 152)). ****P<0.0001.
(C) The bar plots show H3K27me3, H3K27ac, and Lamin B1 corrected total cell fluorescence in HCT116^LMN(B1&B2)-AID^ cells for the control (untreated) and 24-hour auxin treatment conditions. Data was obtained from three technical replicates for each condition (N=3, Control (*n* = 383), 24-hour auxin (*n* = 367)). ****P<0.0001.
(D) Doxycycline treatment alone to induce OsTIR1 expression does not alter *D* in HCT116^LMN(B1&B2)-AID^ cells. N = 3; data was obtained from three independent biological replicates (Untreated (Control) *n* = 507; 24-hr Auxin *n* = 519). ****P<0.0001.
(E) The coefficient of variation plot for HCT116^LMN(B1)-AID^ cells demonstrate chromatin decompaction upon 24 hours of auxin treatment. The truncated violin plots extend from the minimum to the maximum value. The line in the middle of each plot is the median value of the distribution, and the lines above and below are the third and first quartiles, respectively. Data was obtained from three technical replicates for each condition (N=3, Control (*n* = 770), 24-hour auxin (*n* = 492)). ****P<0.0001. Welch’s correction was applied.
(F) The coefficient of variation plot for HCT116^LMN(B2)-AID^ cells demonstrate chromatin decompaction upon 24 hours of auxin treatment. The truncated violin plots extend from the minimum to the maximum value. The line in the middle of each plot is the median value of the distribution, and the lines above and below are the third and first quartiles, respectively. Data was obtained from three technical replicates for each condition (N=3, Control (*n* = 650), 24-hour auxin (*n* = 363)). ****P<0.0001. Welch’s correction was applied.

**Supplementary Figure 6:**

**Degradation of B-type lamins promotes differential gene expression near LAD boundaries.**

(A) Schematic illustration of the three auxin treatment conditions (12 hours, 24 hours, and 24-hour treatment with 6 days of auxin removal) for RNA-sequencing.
(B) Volcano plot showing the top differentially expressed genes (DEGs) for 12 hours of auxin treatment, 48 hours of auxin treatment, and 6 days of auxin withdrawal (adjusted P value < 0.01 and absolute log fold change > 1).
(C) Bar plot showing the number of all DEGs compared to the number of DEGs within versus outside of LAD boundaries defined by publicly available DamID data for all three auxin treatment conditions (adjusted P value < 0.01 and absolute log fold change > 1).
(D) Dot plot showing the ratios of DEGs within LADs, outside of LADs, and all DEGs.
(E) Box plot showing the absolute log fold change and standard error for DEGs within LADs and outside of LADs for all three conditions.
(F) Dot plot showing the Gini coefficient as a measure of transcriptional divergence for all three conditions.

**SUPPLEMENTARY TABLES (submitted as Excel Files)**

**Supplementary Table 1: Plasmids for making parental HCT116 cells expressing OsTIR1.**

| **Plasmid name** | **Transgene** | **Marker** |
| --- | --- | --- |
| pMK232 | CMV-OsTIR1 | Puro |
| pMK364 | CMV-OsTIR1 | loxP-Puro-loxP |
| AAVS1 T2 CRISPR in pX330 | spCas9, AAVS1 T2 gRNA |  |

**Supplementary Table 2: Plasmids for the construction of CRISPR and donor plasmids for tagging.**

| **Plasmid name** | **Transgene** | **Marker** |
| --- | --- | --- |
| pMK290 | mAID-mClover | Hygro |
| AAVS1 T2 CRISPR in pX330 | spCas9, AAVS1 LMNB1 or B2 gRNAs |  |

**Supplementary Table 3: Primers used to generate the sgRNAs for creating the cell lines.**

| **Primer** | **Sequence** |
| --- | --- |
| U6 Promoter Primer 1 | ACTATCATATGCTTACCGTAAC |
| U6 Promoter Primer 2 | gagggcctatttcccatgattc |
| U6 Promoter Primer 3 | aggctgttagagagataattgg |
| LMNB1 CRISPR | GAAGGCTCTGCACTGTATAC |
| LMNB2 CRISPR | GACCCGAGGACCACCTCAAG |
| LMNB1 gRNA Scaffold | gttttagagctaGAAAtagcaagttaaaataaggctagtccgttatcaacttgaaaaagtggcaccgagtcggtgc |
| LMNB2 gRNA Scaffold | gttttagagctaGAAAtagcaagttaaaataaggctagtccgttatcaacttgaaaaagtggcaccgagtcggtgc |
| mClover(A206K) SDM_F | ctgagccatcagtccAAGctgagcaaagacccc |
| mClover(A206K) SDM_R | ggggtctttgctcagCTTggactgatggctcag |
| Linker_Clover_5'_BamHI_SacI | atgcGAGCTCggatccGGTGCAGGCGccgctagcatggtgagcaagggcgaggag |
| Clover_3'_TAA_XbaI | atgctctagattacttgtacagctcgtccatgc |

**Supplementary Table 4: CRISPR-Sirius labeling.**

| **Start** | **End** | **Target Sequences** | **Copies** | **Genes** | **Labeling** |
| --- | --- | --- | --- | --- | --- |
| 1627737 | 1629139 | TGAGGCGGGAAGGGG | 36 | TCF3 | TCF3 |
| 195199025 | 195233876 | ATGATATCACAGTGG | 333 | XXYLT1 | XXYLT1 |

**Supplementary Table 5: Differentially expressed genes for each RNA-seq condition are attached.**
