## Supplementary material for "Depletion of lamins B1 and B2 alters chromatin mobility and induces differential gene expression by a mesoscale-motion dependent mechanism": Key Resources Table

**METHOD DETAILS**

**KEY RESOURCES TABLE**

| REAGENT or RESOURCE | SOURCE | IDENTIFIER |
| --- | --- | --- |
| Bacterial and virus strains | | |
| Lentiviral packaging plasmid pCMV-VSV-G | Addgene | 8454 |
| Lentiviral packaging plasmid pCMV-dR8.2 dvpr | Addgene | 8455 |
| pMK232 (CMV-OsTIR1-PURO) | Addgene | 72834 |
| pMK364 (CMV-OsTIR1-loxP-PURO-loxP) | Addgene | 121184 |
| pX330-U6-Chimeric_BB-CBh-hSpCas9 | Addgene | 42230 |
| AAVS1 T2 CRISPR in pX330 | Addgene | 72833 |
| pMK290 (mAID-mClover-Hygro) | Addgene | 72828 |
| Chemicals, Peptides, and Recombinant Proteins | | |
| Taq DNA Polymerase | G Biosciences | 786-447 |
| Paraformaldehyde 16% solution | Electron Microscopy Sciences | 15710 |
| Triton X-100 | Sigma-Aldrich | 93443 |
| Tween 20 | Sigma-Aldrich | P9416 |
| RIPA Buffer | Sigma-Aldrich | R0278 |
| Protease Inhibitor Cocktail | Sigma-Aldrich | P8340 |
| Sodium Dodecyl Sulfate (SDS) | Sigma-Aldrich | 436143 |
| PBS | Gibco | 10010031 |
| DPBS | Gibco | 14190-144 |
| BSA | Sigma-Aldrich | A7906 |
| EDTA | Thermo Fisher Scientific | 1860851 |
| Igepal CA-630 | Sigma-Aldrich | I3021 |
| MboI Restriction Enzyme | New England Biolabs | R0147 |
| 10X NEBuffer 2 | New England Biolabs | B7002S |
| Biotin-14-dATP | Thermo Fisher Scientific | 19524016 |
| dNTP Solution Mix | New England Biolabs | N0447 |
| T4 PNK | New England Biolabs | M0201 |
| T4 DNA Polymerase 1 | New England Biolabs | M0203 |
| DNA Polymerase 1, Large (Klenow) Fragment | New England Biolabs | M0210 |
| Klenow Exo Minus | New England Biolabs | M0212 |
| 10X NEB T4 DNA Ligase Buffer | New England Biolabs | B0202 |
| 5X NEBNext Quick Ligation Reaction Buffer | New England Biolabs | B6058 |
| DNA Quick Ligase | New England Biolabs | M2200 |
| T4 DNA Ligase | New England Biolabs | M0202 |
| Proteinase K | New England Biolabs | P8102 |
| AMPure XP Beads | Beckman Coulter | A63881 |
| Dynabeads MyOne Streptavidin T1 | Thermo Fisher Scientific | 65601 |
| 3-Indoleacetic acid | Sigma-Aldrich | 12886 |
| Indole-3-acetic acid sodium salt | Sigma-Aldrich | 6505-45-9 |
| Doxycycline Hydrochloride | Fisher Scientific | 10592-13-9 |
| FuGENE HD | Promega | E2311 |
| FuGENE 6 Transfection Reagent | Promega | E2691 |
| Polybrene | Sigma-Aldrich | TR-1003 |
| DAPI Solution | Thermo Fisher Scientific | 62248 |
| Protein Assay Dye Concentrate | BioRad | 500–0006 |
| Blotting Grade Blocker Non-Fat Dry Milk | BioRad | 170–6404 |
| Antibodies | | |
| Alpha Tubulin antibody | Thermo Fisher Scientific | 62204 |
| Anti-Lamin B1 antibody | Cell Signaling | 13435 |
| Anti-Lamin B1 antibody - Nuclear Envelope Marker | Abcam | ab16048 |
| Anti-Lamin B2 antibody | Abcam | ab155319 |
| Anti-Lamin B2 antibody | Cell Signaling | 12255 |
| Invitrogen Goat anti-Rabbit IgG (H+L) Highly Cross-Adsorbed Secondary Antibody, Alexa Fluor 647 | Thermo Fisher Scientific | A-21245 |
| Anti-Rabbit IgG HRP | Promega | W4018 |
| Pierce ECL Western Blotting Substrate | Thermo Fisher Scientific | 32106 |
| Critical Commercial Assays | | |
| Annexin V APC Assay Kit | Cayman Chemical | 601410 |
| Universal Mycoplasma Detection Kit | ATCC | 30-1012K |
| QIAprep Spin Miniprep Kit | Qiagen | 27104 |
| RNeasy Plus Mini Kit | Qiagen | 74134 |
| Applied Biosystems High-Capacity RNA-to-cDNA Kit | Thermo Fisher Scientific | 4387406 |
| NEBNext Ultra Directional RNA Library Prep Kit for Illumina | New England Biolabs | E7760 |
| Qubit 1X dsDNA HS Assay Kit | Thermo Fisher Scientific | Q33230 |
| Experimental Models: Cell lines | | |
| HEK293T | ATCC | CRL-1573 |
| HCT116 | ATCC | CCL-247 |
| Recombinant DNA | | |
| pHAGE-TO-dCas9-P2A-HSA | Addgene | 121938 |
| pHAGE-EFS-MCP-HALOnls | Addgene | 121937 |
| pPUR-hU6-sgRNA-Sirius-8xMS2 | Addgene | 121942 |
| Software and Algorithms | | |
| MATLAB | MathWorks | https://www.mathworks.comproducts/matlab.html |
| ImageJ | NIH | https://imagej.nih.gov/ij/ |
| IDT Custom Alt-R Guide Design | IDT | https://sg.idtdna.com/site/order/designtool/index/CRISPR_CUSTOM |
| WEG CRISPR finder | Sanger Institute | https://www.sanger.ac.uk/htgt/wge/ |
| Snap Gene | Insightful Science | Snapgene.com |
| Juicebox | Robinson et al., 2018 | https://www.aidenlab.org/juicebox/ |
| Python | Python Software Foundation | http://www.python.org |
| GraphPad Prism | GraphPad Software, Inc. | https://graphpad.com |
| R | R Core Team 2022 | https://www.R-project.org/ |
| Other | | |
| McCoy’s 5A Modified Medium | Thermo Fisher Scientific | 16600-082 |
| Dulbecco’s Modified Eagle’s Medium | Gibco | 11965-092 |
| Fetal bovine serum | Thermo Fisher Scientific | 16000-044 |
| Penicillin-streptomycin | Thermo Fisher Scientific | 31985062 |
| Hygromycin B Gold | Gibco | 10687010 |
| Opti-MEM I Reduced Serum Medium | Gibco | 31985070 |
| McCoy’s 5A Medium | Cytiva | SH30270.01 |
| 0.25% Trypsin-EDTA | Gibco | 25200-056 |
| Invitrogen UltraPure DNase/ RNase-Free Distilled Water | Fisher Scientific | 10-977-015 |
| Janelia Fluor 646 HaloTag | Promega | GA1120 |
| 6-well glass bottom plates | Cellvis | P06-1.5H-N |
| 12-well glass bottom plates | Cellvis | P12-1.5H-N |
| 24-well glass bottom plates | Cellvis | P24-1.5H-N |
| Corning tissue-culture treated cell culture dishes (10-cm plates) | Millipore Sigma | CLS430165 |
| Millex-HV Syringe Filter Unit, 0.45 µm, PVDF, 33 mm, gamma sterilized | Millipore Sigma | SLHVR33RS |
